# Flotillin-2 dampens T cell antigen-sensitivity and functionality

**DOI:** 10.1101/2024.04.26.591344

**Authors:** Sookjin Moon, Fei Zhao, Mohammad N. Uddin, Charles J. Tucker, Peer W. F. Karmaus, Michael B. Fessler

## Abstract

T cell receptor (TCR) engagement triggers T cell responses, yet how TCR-mediated activation is regulated at the plasma membrane remains unclear. Here, we report that deleting the membrane scaffolding protein Flotillin-2 (Flot2) increases T cell antigen sensitivity, resulting in enhanced TCR signaling and effector function to weak TCR stimulation. T cell-specific Flot2-deficient mice exhibited reduced tumor growth and enhanced immunity to infection. Flot2-null CD4^+^ T cells exhibited increased T helper 1 polarization, proliferation, Nur77 induction, and phosphorylation of ZAP70 and LCK upon weak TCR stimulation, indicating a sensitized TCR-triggering threshold. Single cell-RNA sequencing suggested that Flot2*-*null CD4^+^ T cells follow a similar route of activation as wild-type CD4^+^ T cells but exhibit higher occupancy of a discrete activation state under weak TCR stimulation. Given prior reports that TCR clustering influences sensitivity of T cells to stimuli, we evaluated TCR distribution with super-resolution microscopy. Flot2 ablation increased the number of surface TCR nanoclusters on naïve CD4^+^ T cells. Collectively, we posit that Flot2 modulates T cell functionality to weak TCR stimulation, at least in part, by regulating surface TCR clustering. Our findings have implications for improving T cell reactivity in diseases with poor antigenicity, such as cancer and chronic infections.

## Introduction

T cells encounter a vast array of antigens, but few cognate antigens induce T cell receptor (TCR) triggering and T cell activation. Proper regulation of TCR triggering plays a critical role in protective T cell immunity, but how T cells regulate TCR triggering remains unclear. The kinetic proofreading model, introduced in 1995, proposed that the duration of TCR interaction with the peptide-major histocompatibility complex (pMHC) is a critical determinant of TCR discrimination (1–6). By contrast, several studies have more recently suggested spatial organization of receptors on the membrane as a key determinant of TCR activation. Thus, in the engineered TCR signaling system, prolonged interaction alone is not sufficient to initiate TCR signaling; instead, receptor clustering on the membrane is necessary (7). Moreover, signal strength reportedly amplifies as the inter-ligand spacing decreases (8) and the density of TCR-CD3 complexes within nanoclusters determines TCR triggering efficiency (9), together highlighting the importance of spatial thresholds in TCR triggering. Despite the accumulating evidence highlighting the role of surface TCR clustering in the initiation of TCR signaling, the regulation of TCR spatial organization on the membrane and its impact on fine-tuning of the TCR triggering threshold has remained elusive.

Flotillin-1 (Flot1) and flotillin-2 (Flot2) are evolutionarily conserved, ubiquitously expressed scaffolding proteins (10–15) that are thought to localize to lipid rafts, membrane microdomains that support receptor-mediated signaling (16–24). Rafts have been suggested to promote TCR signaling by supporting assembly of the immunological synapse and/or assisting signaling by costimulatory molecules (25, 26). Indeed, Flot2 has recently been implicated in TCRζ trafficking, but divergent findings have been reported on its role in TCR signaling perhaps in part because of reliance on Jurkat T cell lines (27, 28). Overall, these conflicting observations underscore the need for further elucidation of the role of flotillins in T cell responses and their potential for therapeutic intervention.

Here, we investigated the role of Flot2 in TCR triggering and T cell responses using both *in vivo* and *in vitro* models. We found that Flot2-deficient mice exhibited delayed tumor growth and heightened resistance to *Listeria* infection, associated with augmented effector T cell proliferation and cytokine production. *In vitro* models revealed that Flot2-deficient CD4^+^ T cells exhibited enhanced TCR activation, even under weak TCR stimulation. Furthermore, Flot2-deficient CD4^+^ T cells showed heightened differentiation toward a T helper (Th)1 subset upon exposure to various stimulus concentrations, including weak TCR stimulation. This enhanced differentiation is likely due to a sensitized TCR triggering threshold, as suggested by elevated phosphorylation of signaling proteins in the proximal TCR signaling pathway following weak stimulation. Single-cell RNA sequencing (scRNA-seq) analysis suggested that *Flot2^CD4^* CD4^+^ T cells follow a similar route of activation as *Flot2^WT^* CD4^+^ T cells but exhibit higher occupancy in a discrete activation state under weak TCR stimulation. Finally, super-resolution imaging revealed an increased number of surface TCR nanoclusters on Flot2-deficient naïve CD4^+^ T cells, suggesting that Flot2 controls spatial organization of TCR molecules in the steady-state and thereby sensitivity of naïve T cells to the environment.

## Results

### Flot2 deletion enhances anti-tumor activity of T cells in murine tumor models

To investigate the role of Flot2 in T cell responses, we generated *Flot2* global knockout mice (i.e., *Flot2^−/−^* mice) by flanking the *Flot2* coiled-coil domain (14) with loxP sites and then crossing these *Flot2^flox/flox^*mice with CMV-Cre mice (29) (Supplementary Figure 1). B16F10 melanoma and MC38 colon adenocarcinoma, well-established *in vivo* models of T cell anti-tumor immunity (30–33), were first tested. In both the B16F10 and MC38 models, *Flot2^−/−^* mice exhibited delayed/reduced tumor growth compared to *Flot2*^+/+^ counterparts (Figure 1a,b). Similar results were noted in mice of both sexes. Furthermore, *Flot2^−/−^* mice exhibited an increased frequency of CD4^+^ and CD8^+^ tumor-infiltrating lymphocytes (TILs) in the B16F10 melanoma model, specifically including Ki67^+^ proliferating effector CD4^+^ and CD8^+^ cells (Figure 1c-j). The expression of TOX, a marker of functional exhaustion, was decreased in CD8^+^ TILs of *Flot2^−/−^* mice with MC38 tumors, whereas TIM-3 expression was similar to that in wild type controls (Supplementary Figure 2a-f). Total splenic IFN-γ production to melanoma TRP-2 peptide stimulation was also elevated in B16F10 tumor-bearing *Flot2^−/−^* mice compared to their *Flot2^+/+^* counterparts, indicating an enhanced response to tumor antigen (Figure 1k). Taken together, these results indicated an augmented anti-tumor immune response in Flot2-deficient mice.

**Figure 1.**
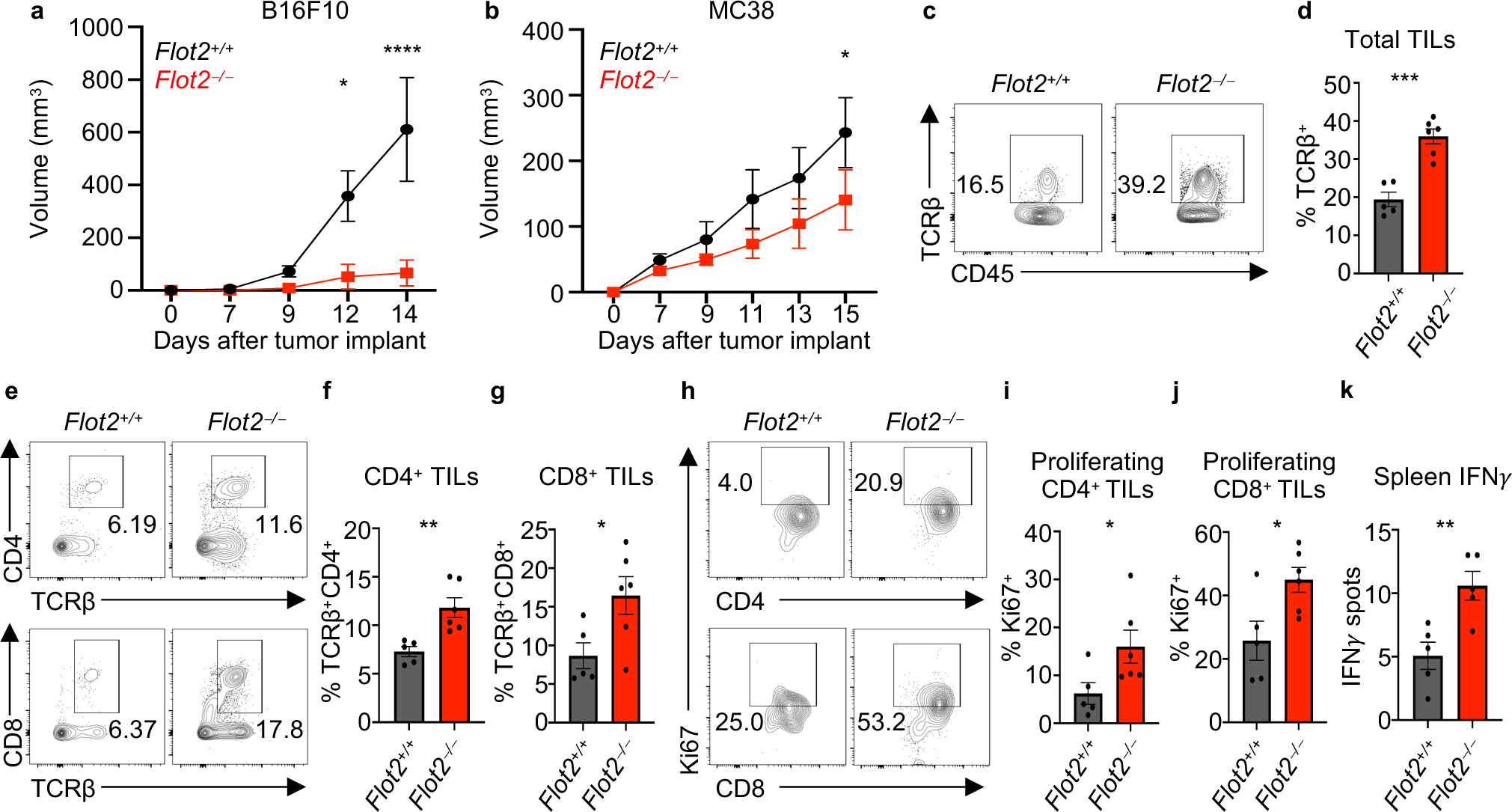
Flot2 deficiency potentiates the anti-tumor activity of both CD4^+^ and CD8^+^ T cells *in vivo*. **a, b** Tumor volume in *Flot2^+/+^* or *Flot2^−/−^* mice injected with B16F10 (**a**) or MC38 (**b**) (n = 7 per group). **c-j** Flow cytometric analysis of tumor-infiltrating lymphocytes (TILs) in B16F10 tumor-bearing *Flot2^+/+^* or *Flot2^−/−^* mice: Representative plots (**c**, **e**, **h**) are shown. TCRβ^+^ (**d**), TCRβ^+^CD4^+^ (**f**), and TCRβ^+^CD8^+^ (**g**) populations within 7AAD^−^CD45^+^ population, and Ki67^+^ populations among 7AAD^−^CD45^+^TCRβ^+^CD4^+^CD44^+^CD62L^−^ population (**i**) or 7AAD^−^CD45^+^TCRβ^+^CD8^+^CD44^+^CD62L^−^ population (**j**) are depicted. **k** Splenocytes from B16F10 tumor-bearing *Flot2^+/+^* or *Flot2^−/−^* mice were stimulated with 1 μg/ml of TRP-2 melanoma peptide for 24 hours and assayed for antigen-specific reactivity using an IFNγ ELISpot assay. Data are representative of two independent experiments. Data were analyzed by unpaired t-test (**d**, **f**, **g**, **i-k**) or two-way ANOVA followed with Sidak’s multiple comparison tests (**a**, **b**). Error bars denote SEM; *P<0.05; **P<0.01; ***P<0.001; ****P<0.0001.

Next, we sought to examine whether Flot2-deficiency specifically within the T cell compartment is sufficient to confer enhanced anti-tumor immunity. To explore this, we generated mixed bone marrow chimeras by reconstituting lethally irradiated TCRα-deficient (*TCRα^−/−^*) recipient mice with a 1:5 ratio mixture of bone marrow cells from either *Flot2^+/+^*or *Flot2^−/−^* donor mice and *TCRα^−/−^* mice, thereby generating mice with predominantly wild-type hematopoietic cells, except for a Flot2-deficient T cell compartment (34). Subsequently, these chimeras were inoculated with B16F10 tumors. Notably, *TCRα^−/−^* recipients reconstituted with *Flot2^−/−^* bone marrow exhibited a significant reduction in tumor volumes compared to those reconstituted with WT bone marrow (Supplementary Figure 2g). Furthermore, we observed increased proliferation marker Ki67 expression in both CD4^+^ and CD8^+^ TILs in the chimeras transferred with *Flot2^−/−^* bone marrow, along with an expansion of the CD44^+^IFNγ^+^ population within CD8^+^ TILs (Supplementary Figure 2h-j). These findings suggest that Flot2 deficiency specifically within T cells augments anti-cancer immune responses.

This finding prompted us to generate T cell-specific Flot2-deficient mice through crossbreeding of *Flot2^flox/flox^* mice with *CD4^Cre^* mice (i.e., *Flot2^CD4^* mice) in order to further explore the T cell-intrinsic role of Flot2 in anti-cancer immunity (Supplementary Figure 3a-c). Flot2 deletion in T cells did not cause overt abnormalities in thymocyte development (Supplementary Figure 3d-h). In the steady state, peripheral T cells in the lymph nodes of *Flot2^CD4^*mice exhibited similar characteristics to those in *Flot2^WT^* mice. There was, however, a marginal decrease in total CD4^+^ T cell numbers, accompanied by an increase in the percentage of the CD44^+^CD62L^-^ population, as well as enhanced expression of Nur77, T-bet, and LFA-1α within CD4^+^ T cells, suggesting a shift from naïve to activated status (Supplementary Figure 3i-n).

As above for *Flot2^−/−^* mice, *Flot2^CD4^* mice were evaluated in the B16F10 melanoma and MC38 colon adenocarcinoma models. Consistent with the phenotypes of *Flot2^−/−^* mice, *Flot2^CD4^* mice showed reduced growth of both B16F10 and MC38 tumors compared to *Flot2^WT^* controls (Figure 2a,b). *Flot2^CD4^* mice also showed elevated populations of CD4^+^ and CD8^+^ TILs, as well as increased Ki67^+^ proliferating effector CD4^+^ and CD8^+^ cells in the B16F10 melanoma model (Figure 2c-h). Moreover, we observed heightened expression of IFNγ and TNFα effector cytokines in CD4^+^ T cells within the tumor-draining lymph nodes (dLN) of *Flot2^CD4^* mice (Figure 2i-k). However, no discernible difference was observed in CD8^+^ T cells within the dLN (Figure 2l-n). Overall, these data demonstrate that specific Flot2-deficiency in T cells boosts anti-tumor responses of both CD4^+^ and CD8^+^ T cells in murine tumor models.

**Figure 2.**
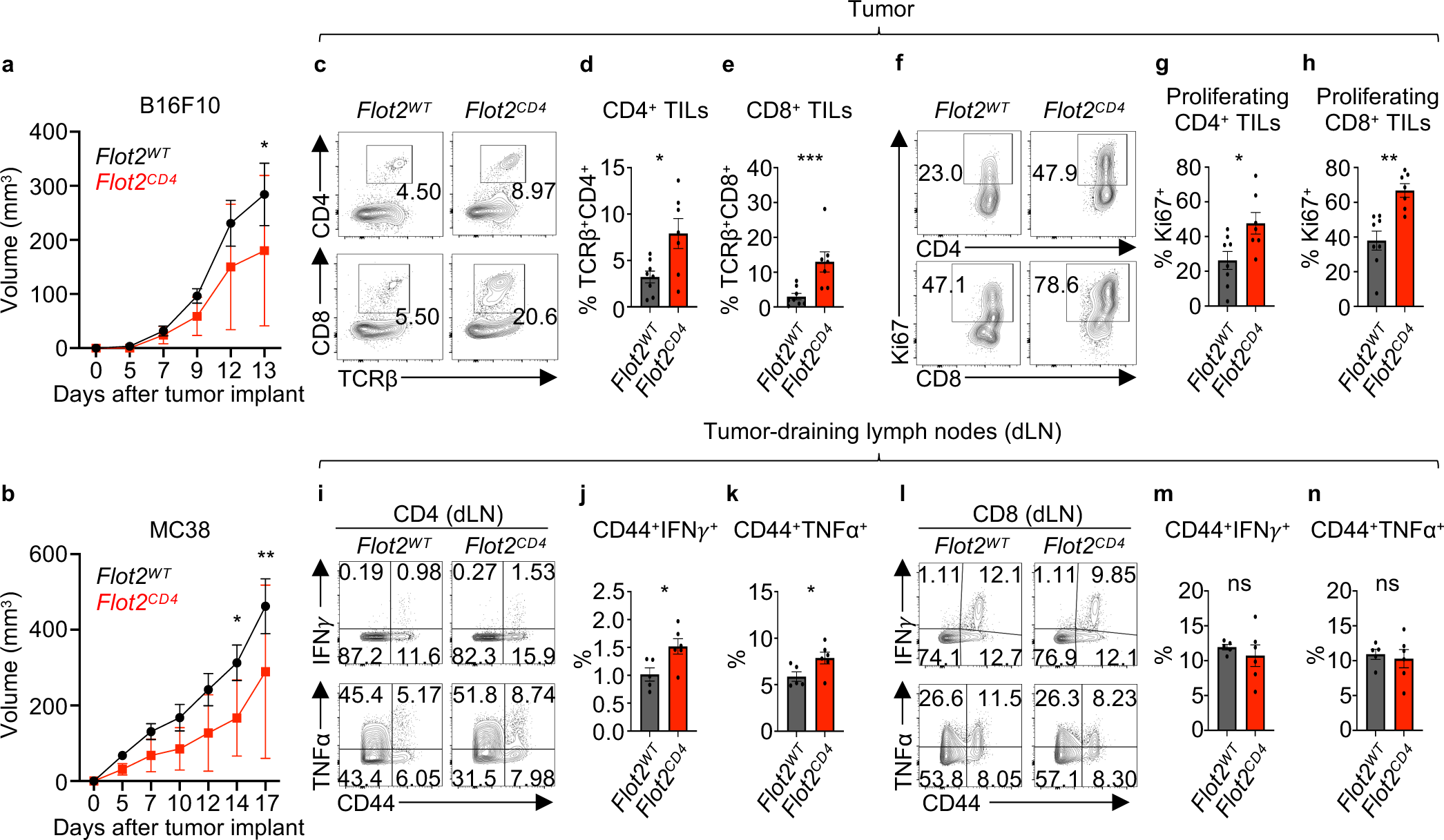
T cell-specific Flot2 deficiency potentiates the anti-tumor activity of both CD4^+^ and CD8^+^ T cells *in vivo*. **a, b** Tumor volume in *Flot2^WT^* or *Flot2^CD4^* mice injected with B16F10 (**a**; n = 10 per group) or MC38 (**b**, n = 9 for *Flot2^WT^* and n = 14 for *Flot2^CD4^*). **c**-**h** Flow cytometric analysis of TILs: Representative plots (**c**, **f**) are shown. TCRβ^+^CD4^+^ (**d**) and TCRβ^+^CD8^+^ (**e**) populations among 7AAD^−^CD45.2^+^ population, and Ki67^+^ populations among 7AAD^−^CD45.2^+^TCRβ^+^CD4^+^ population (**g**) or 7AAD^−^CD45.2^+^TCRβ^+^CD8^+^ population (**h**) in B16F10-bearing *Flot2^WT^* or *Flot2^CD4^* mice are presented. **i**-**n** Flow cytometric analysis of tumor-draining lymph nodes (dLNs): Representative plots (**i**, **l**) are displayed. CD44^+^IFNγ^+^ (**j**) and CD44^+^TNFα^+^ (**k**) populations among 7AAD^−^CD45.2^+^TCRβ^+^CD4^+^ population and CD44^+^IFNγ^+^ (**m**) and CD44^+^TNFα^+^ (**n**) among 7AAD^−^CD45.2^+^TCRβ^+^CD8^+^ population in B16F10-bearing *Flot2^WT^* or *Flot2^CD4^* mice are indicated. Data are representative of two independent experiments (**a**-**n**). Data were analyzed by unpaired t-test (**d**, **e**, **g**, **h**, **j**, **k**, **m**, **n**) or two-way ANOVA followed with Sidak’s multiple comparison tests (**a**, **b**). Error bars denote SEM; *P<0.05; **P<0.01; ***P<0.001. ns = non-significant.

### Flot2 deficiency boosts the anti-bacterial responses of CD4^+^ and CD8^+^ T cells *in vivo*

To evaluate the role of Flot2 in regulating anti-bacterial immune responses *in vivo*, we next challenged mice with *Listeria monocytogenes*. Following infection, *Flot2^−/−^* mice showed elevated resistance to weight loss compared to *Flot2^+/+^* mice (Figure 3a). Furthermore, both CD4^+^ and CD8^+^ T cells in *Flot2^−/−^* mice exhibited increased expression of Ki67 and TNFα compared to *Flot2^+/+^* mice (Figure 3b-g), suggesting increased proliferation and effector function. Consistent with this, CD44^+^T-bet^+^, CD44^+^IFNγ^+^, CD44^−^TNFα^+^, and CD44^+^IL-2^+^ populations were all augmented in both CD4^+^ and CD8^+^ splenic T cells (Supplementary Figure 4). Given these findings, we next used *Flot2^CD4^* mice to investigate whether T cell-specific Flot2-deficiency also improves the anti-bacterial T cell response. Notably, *Flot2^CD4^* mice also showed less weight loss compared to *Flot2^WT^* after *L. monocytogenes* infection (Figure 3h).

**Figure 3.**
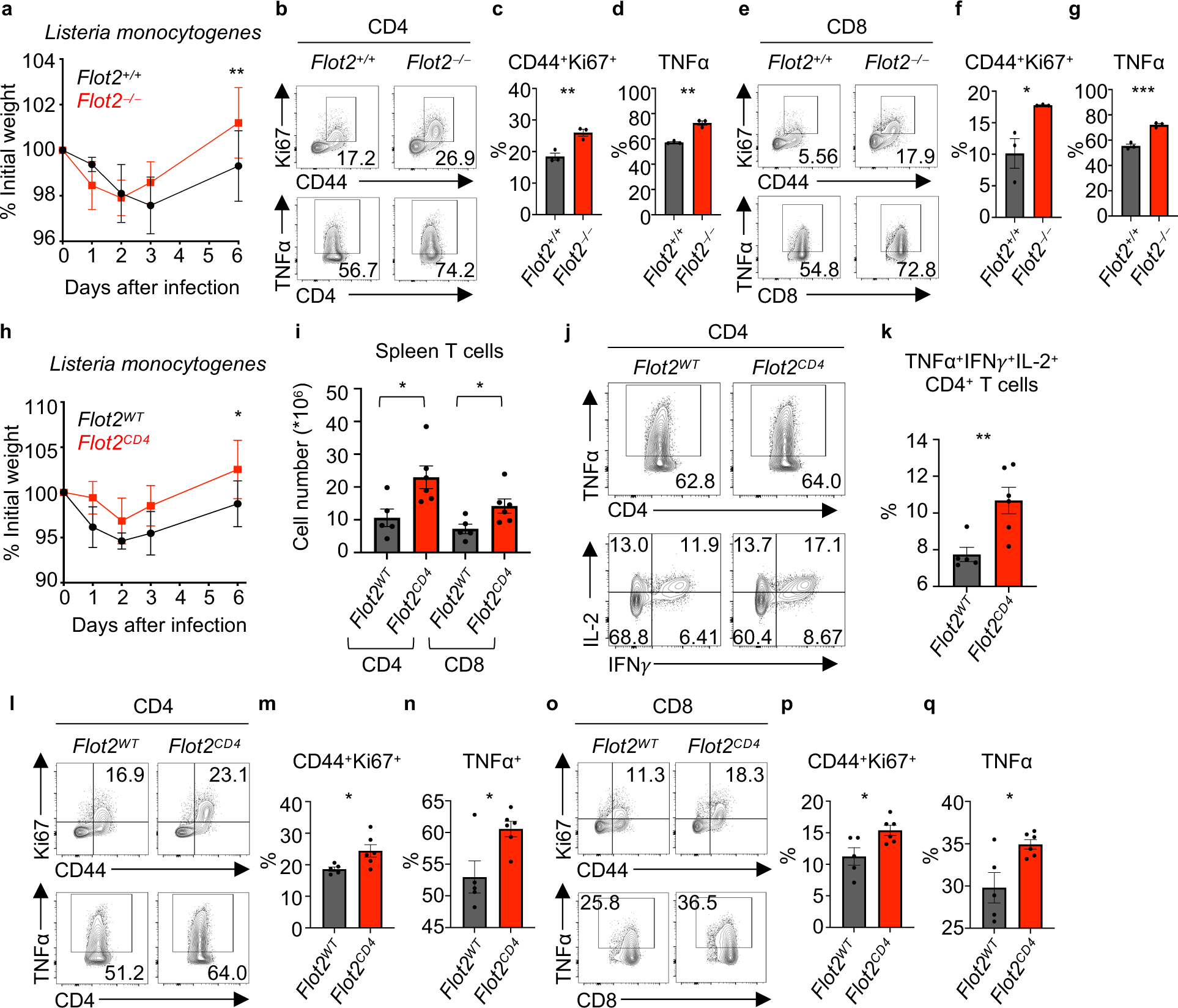
Flot2 deficiency promotes CD4^+^ and CD8^+^ T cell responses against *Listeria monocytogenes* infection. **a** Weight loss in *Flot2^+/+^* or *Flot2^−/−^* mice following *Listeria monocytogenes* infection (5000 CFU per mouse) is presented as the mean percentage of initial weight (n = 4-5 per group). **b**-**g** Flow cytometric analysis of splenic T cells from *Listeria*-infected *Flot2^+/+^* or *Flot2^−/−^* mice: Representative plots (**b**, **e**) are provided. CD44^+^Ki67^+^ (**c**) and TNFα^+^ (**d**) populations within viable CD45^+^TCRβ^+^CD4^+^ population, and CD44^+^Ki67^+^ (**f**) and TNFα^+^ (**g**) populations within viable CD45^+^TCRβ^+^CD8^+^ population are shown. **h** Weight loss in *Flot2^WT^* or *Flot2^CD4^* mice following *Listeria monocytogenes* infection (5000 CFU per mouse) is presented as the mean percentage of initial weight (n = 5-6 per group). **i** Splenic CD4^+^ and CD8^+^ T cells numbers in infected *Flot2^WT^* or *Flot2^CD4^* mice are depicted. **j-q** Flow cytometric analysis of splenic T cells from *Listeria*-infected *Flot2^WT^* or *Flot2^CD4^* mice: Representative plots (**j**, **l, o**) are shown. TNFα^+^IFNγ^+^IL-2^+^ (**k**), CD44^+^Ki67^+^ (**m**), TNFα^+^ (**n**) populations within viable CD45^+^TCRβ^+^CD4^+^ population, and CD44^+^Ki67^+^ (**p**) and TNFα^+^ (**q**) populations within viable CD45^+^TCRβ^+^CD8^+^ population are shown. Data are representative of two independent experiments (**a**-**q**). Data were analyzed by unpaired t-test (**c**, **d**, **f, g, i**, **k**, **m**, **n**, **p**, **q**) or two-way ANOVA followed with Sidak’s multiple comparison tests (**a**, **h**). Error bars denote SEM; *P<0.05; **P<0.01; ***P<0.001.

Quantification of the absolute number of T cells in the spleen of infected *Flot2^WT^* and *Flot2^CD4^* mice revealed an increase in both CD4^+^ and CD8^+^ T cell numbers in *Flot2^CD4^*mice (Figure 3i). Moreover, *Flot2^CD4^* mice had increased TNFα^+^IFNγ^+^IL-2^+^ multifunctional CD4^+^ T cells, which play a crucial role in infection control (35, 36) (Figure 3j,k). Consistent with findings in *Flot2^−/−^* mice, *Flot2^CD4^*mice also exhibited heightened expression of Ki67 and TNFα in both CD4^+^ and CD8^+^ splenic T cells (Figure 3l-q). These data collectively suggest that Flot2 deficiency improves the responses of both CD4^+^ and CD8^+^ T cells in the context of *in vivo* infection with *L. monocytogenes*.

### TCR activation induces enhanced response in Flot2-deficient CD4^+^ but not CD8^+^ T cells

After confirming that Flot2 deficiency enhances effector T cell responses *in vivo* in both tumor and infection models, we investigated this phenomenon mechanistically, using reductionist *in vitro* approaches. Purified naïve CD4^+^ and CD8^+^ T cells from *Flot2^WT^*or *Flot2^CD4^* mice were stimulated with increasing concentrations of plate-bound anti-CD3, along with a fixed concentration of soluble anti-CD28 (1 μg/ml), modeling TCR stimulation and co-stimulation exclusively. Following stimulation, both *Flot2^WT^* and *Flot2^CD4^* CD4^+^ T cells showed a concentration-dependent increase in CellTrace Violet^−^ (CTV^−^; i.e., proliferated), Ki67^+^, T-bet^+^, and CD25^+^ populations, confirming T cell activation in line with the strength of TCR stimulation (Figure 4a-d and Supplementary Figure 5a). *Flot2^CD4^* CD4^+^ T cells exhibited heightened proliferation (CTV^−^ and Ki67^+^) and T-bet and CD25 expression compared to *Flot2^WT^* across various concentrations of anti-CD3 (Figure 4a-d and Supplementary Figure 5a). Remarkably, Flot2 deficiency augmented CD4^+^ T cell proliferation even at very low concentrations of anti-CD3 (0.0625 μg/ml) (Figure 4a) and increased expression of the early T cell activation marker (CD25) at low concentrations of anti-CD3 (0.125 μg/ml) (Figure 4d), highlighting enhanced T cell responsiveness to weak TCR stimulation. By contrast, Flot2-deficient CD8^+^ T cells did not experience increased cell proliferation or early activation compared to WT controls (Figure 4e-h and Supplementary Figure 5b). Collectively, these findings indicate that Flot2 deficiency renders CD4^+^ T cells more responsive to weak TCR stimulation in the presence of both TCR stimulation and co-stimulation, whereas Flot2-deficient CD8^+^ T cells may rely on supplementary factors, potentially available *in vivo*, for their boosted activation.

**Figure 4.**
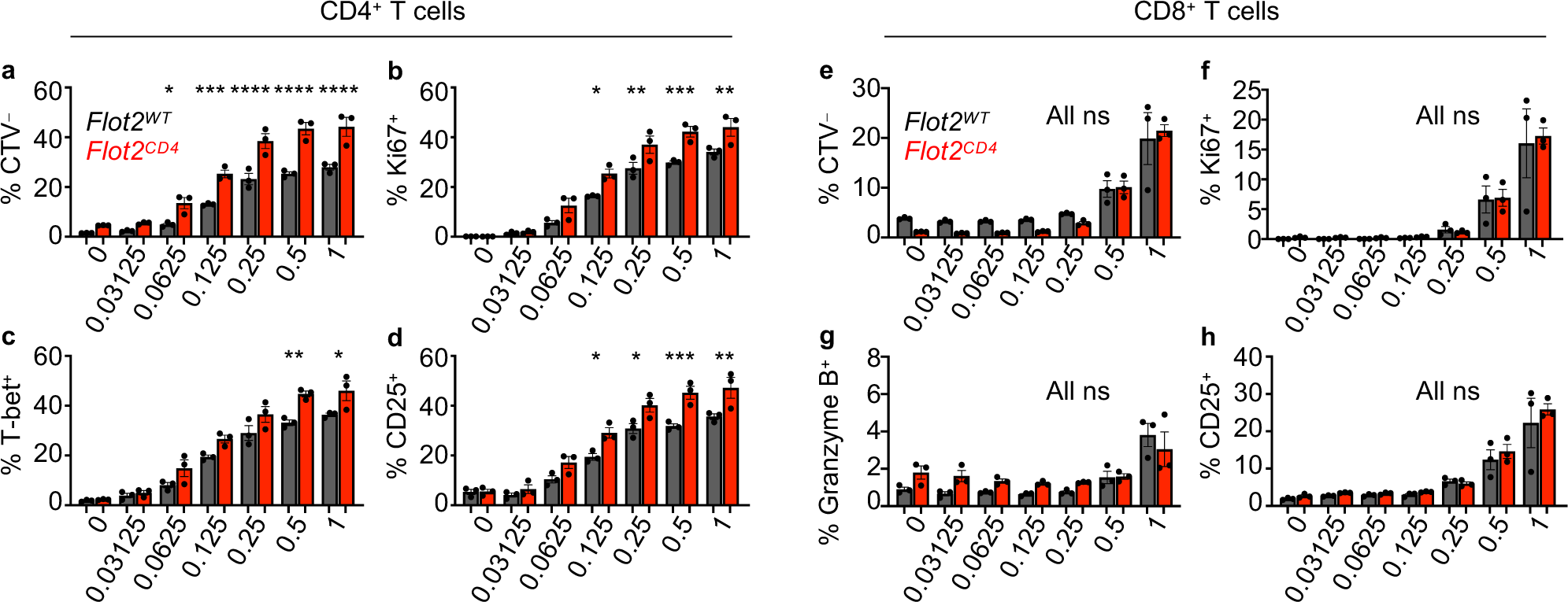
*Flot2^CD4^* CD4^+^ T cells, but not *Flot2^CD4^* CD8^+^ T cells, showed enhanced T cell responses in *in vitro* T cell stimulation. **a-d** Naïve CD4^+^ T cells were purified and stimulated *in vitro* for 72 hours with varying doses of plate-bound anti-CD3, alongside a fixed dose of soluble anti-CD28 (1 μg/ml), followed by flow cytometric analysis to assess cell proliferation and activation. CTV^−^ (**a**), Ki67^+^ (**b**), T-bet^+^ (**c**), and CD25^+^ (**d**) populations within viable TCRβ^+^CD4^+^ population are shown. **e**-**h** Naïve CD8^+^ T cells were purified and stimulated *in vitro* for 72 hours with varying doses of plate-bound anti-CD3, alongside a fixed dose of soluble anti-CD28 (1 μg/ml), followed by flow cytometric analysis to assess cell proliferation and activation. CTV^−^ (**e**), Ki67^+^ (**f**), Granzyme B^+^ (**g**), and CD25^+^ (**h**) populations within viable TCRβ^+^CD8^+^ population are shown. Data are representative of two independent experiments (**a**-**h**). Data were analyzed by one-way ANOVA followed with Sidak’s multiple comparison tests (**a**-**h**). Error bars denote SEM; *P<0.05; **P<0.01; ***P<0.001; ****P<0.0001. ns = non-significant.

### Deletion of Flot2 in CD4^+^ T cells promotes Th1 cell differentiation

During T cell activation, naïve CD4^+^ T cells have the potential to differentiate into various helper T cell subsets, characterized by distinct transcription factors, cytokines, and functions (37–39). The determination of cell fate during differentiation is influenced by both TCR signal strength and the cytokine milieu, with recent findings suggesting an association between strong TCR signals and Th1 differentiation (40–42). Since *in vitro* stimulated Flot2-deficient CD4^+^ T cells express higher levels of CD25 and T-bet (Figure 4c,d), indicative of strong TCR signal strength, we investigated the impact of Flot2 deficiency on T helper differentiation. Initially, naïve CD4^+^ T cells were differentiated *in vitro* using Th1 polarizing medium and varying concentration of plate-bound anti-CD3. Both *Flot2^WT^*and *Flot2^CD4^* CD4^+^ T cells exhibited concentration-dependent induction of Th1 polarization and proliferation (Figure 5). Notably, *Flot2^CD4^* CD4^+^ T cells displayed a significant increase in T-bet^+^IFNγ^+^ and CD44^+^TNFα^+^ populations compared to *Flot2^WT^* CD4^+^ T cells across various concentrations of anti-CD3, even at very low concentrations (Figure 5a-d). Furthermore, Th1 cell proliferation of *Flot2^CD4^* CD4^+^ T cells was also robustly induced at low concentrations of anti-CD3, likely due to their increased sensitivity to TCR stimulation (Figure 5e,f).

**Figure 5.**
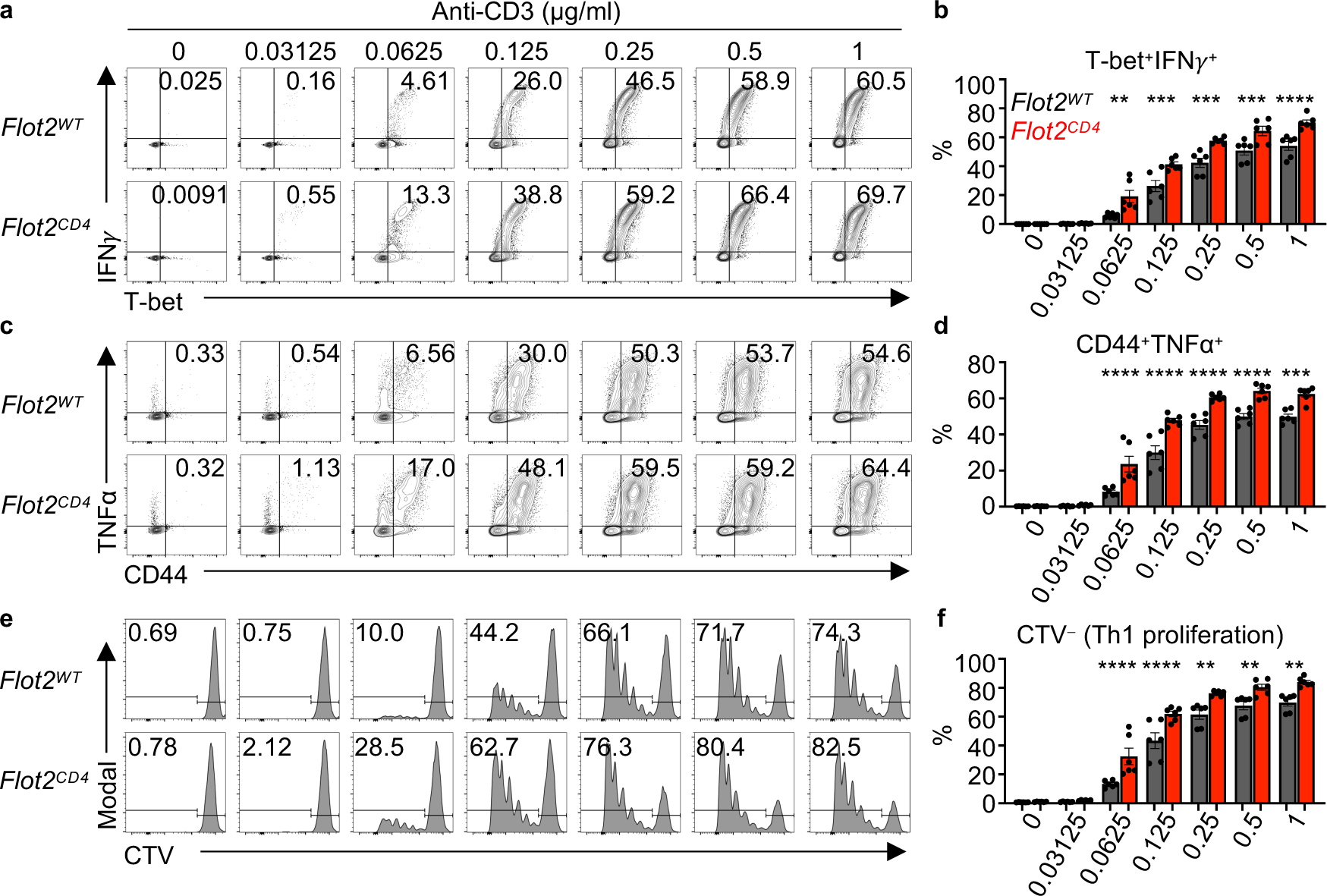
Flot2 ablation promotes CD4^+^ T cell differentiation into T helper 1 upon weak TCR stimulation. **a-f** Naïve CD4^+^ T cells were purified and differentiated toward the T helper (Th) 1 subtype *in vitro* using Th1 polarizing conditions, followed by flow cytometric analysis to assess Th1 polarization, cytokine production, and cell proliferation. Representative plots (**a**, **c, e**) are shown. T-bet^+^IFNγ^+^ (**b**), CD44^+^TNFα^+^ (**d**), and CTV^−^ populations (**f**) within viable TCRβ^+^CD4^+^ population are shown. Data are representative of three independent experiments (**a**-**f**). Data were analyzed by one-way ANOVA followed with Sidak’s multiple comparison tests (**b**, **d**, **f**). Error bars denote SEM; **P<0.01; ***P<0.001; ****P<0.0001.

Next, we evaluated differentiation into other Th subsets using Th2, Th17, or Treg polarizing conditions. In contrast to Th1 differentiation, *Flot2^CD4^*CD4^+^ T cells showed no difference in Th2 proliferation, evidenced by the CTV^−^ population, and a decrease in IL-4^+^GATA3^+^ populations at sufficient TCR stimulation (Supplementary Figure 6a-d). Similarly, no difference was observed in Th17 differentiation (Supplementary Figure 6e,f). However, *Flot2^CD4^* CD4^+^ T cells showed augmented differentiation into Foxp3^+^ Treg populations selectively upon weak TCR stimulation (Supplementary Figure 6g,h). Given that Treg differentiation tends to favor low-abundance, high-affinity antigens(43), this finding may reflect the increased reactivity of *Flot2^CD4^* CD4^+^ T cells to the low abundance of anti-CD3 compared to *Flot2^WT^* CD4^+^ T cells. On the basis of these findings, we conclude that Flot2-deficient CD4^+^ T cells are inclined towards Th1 and Treg differentiation even at very low concentrations of anti-CD3, likely due to their hypersensitivity to weak TCR stimulation.

### Flot2 ablation decreases TCR triggering threshold in CD4^+^ T cells

Given the heightened proliferation, activation, and Th1 differentiation observed in Flot2-deficient CD4^+^ T cells, we hypothesized an augmentation in TCR signaling in CD4^+^ T cells upon Flot2 ablation. To test this, we stimulated purified naïve CD4^+^ T cells from *Flot2^WT^* or *Flot2^CD4^* mice with varying concentrations of plate-bound anti-CD3 *in vitro* and analyzed them after 3 hours or 24 hours to observe early-phase T cell activation. *Flot2^CD4^* CD4^+^ T cells exhibited elevated expression of Nur77, a marker of TCR signal strength, particularly at low concentrations of anti-CD3 (Figure 6a,b). Additionally, the early T cell activation marker CD69 was increased in *Flot2^CD4^*CD4^+^ T cells compared to *Flot2^WT^* CD4^+^ T cells (Figure 6a,c). Conversely, *Flot2^CD4^* CD8^+^ T cells did not exhibit enhanced expression of Nur77 and CD69, further emphasizing the necessity of additional factors for the Flot2-dependent activation phenotype in CD8^+^ T cells (Supplementary Figure 7a-c). We next investigated earlier phases of TCR signaling by profiling phosphorylation of TCR signaling molecules after 3 minutes of *in vitro* stimulation. Notably, anti-CD3/-CD28-induced early phosphorylation of ZAP70 and LCK were enhanced in *Flot2^CD4^* CD4^+^ T cells compared to *Flot2^WT^*CD4^+^ T cells (Figure 6d).

**Figure 6.**
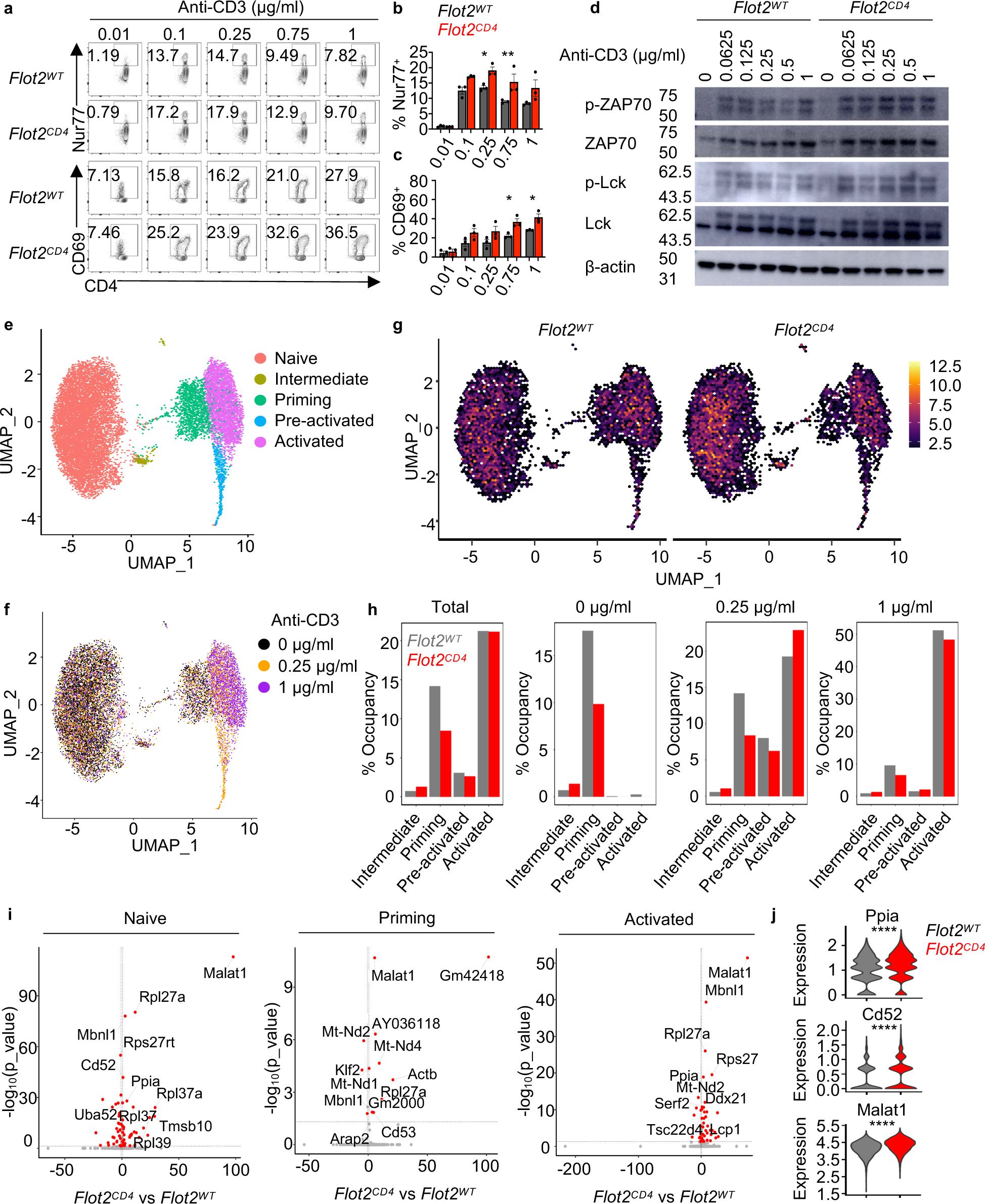
Flot2 ablation sensitizes TCR triggering threshold in CD4^+^ T cells. **a**-**c** Naïve CD4^+^ T cells were purified and stimulated *in vitro* for 3 hours (**b**) or 24 hours (**c**) with varying doses of plate-bound anti-CD3, alongside a fixed dose of soluble anti-CD28 (1 μg/ml), followed by flow cytometric analysis to assess TCR signaling (Nur77) and early T cell activation (CD69). Representative plots (**a**) are shown. Nur77^+^ (**b**) and CD69^+^ (**c**) populations within viable TCRβ^+^CD4^+^ population are shown. **d** Phosphorylation of TCR signaling molecules in naïve CD4^+^ T cells stimulated with varying doses of plate-bound anti-CD3, alongside a fixed dose of soluble anti-CD28 (1 μg/ml), for 3 minutes. **e**, **f** scRNA-seq was performed on naive CD4^+^ T cells following a 3 hour stimulation with varying concentrations (0, 0.25, or 1 μg/ml) of plate-bound anti-CD3. A fixed dose of soluble anti-CD28 (1 μg/ml) was provided under the conditions of 0.25 or 1 μg/ml of plate-bound anti-CD3. Unsupervised T cell clusters were annotated as five distinguishable functional states on a UMAP plot (**e**), and the effect of stimulation dose was assessed by projection onto the UMAP (**f**). **g** Distribution of *Flot2^WT^* or *Flot2^CD4^* CD4^+^ T cells in each cluster. **h** Occupancy of *Flot2^WT^* or *Flot2^CD4^* genotype in each T cell clusters (Intermediate, priming, pre-activated, and activated), analyzed by total or each stimulatory conditions. **i** Volcano plots of spliced RNA in naïve, priming, and activated clusters. **j** Expression level of spliced RNA related to cellular proliferation in the naïve cluster. Data are pooled from three independent experiments (**a**-**c**) or representative of two independent experiments (**d**). Data were analyzed by one-way ANOVA followed with Sidak’s multiple comparison tests (**a**-**c**) or Wilcoxon Rank Sum test (**j**). Error bars denote SEM; *P<0.05; **P<0.01; ****P<0.0001.

To gain a more comprehensive understanding of the role of Flot2 in CD4^+^ T cells during T cell activation, we next performed scRNA-seq on naïve *Flot2^WT^* or *Flot2^CD4^* CD4^+^ T cells after *in vitro* stimulation with varying concentrations of anti-CD3 antibody-mediated TCR stimulation: no (0 μg/ml), weak (0.25 μg/ml), or strong (1 μg/ml) stimulation for 3 hours. Clustering the results using the Leiden algorithm revealed five distinct functional states based on expression of marker genes: naïve, intermediate, priming, pre-activated, and activated as visualized using Uniform Manifold Approximation and Projection (UMAP) (Figure 6e, Supplementary Figure 7d).

These clusters aligned well with previously reported gene sets related to early T cell activation (44) and hallmark genes of T cell activation (*Nr4a1*, *Myc*, *Cd69*, *Il2rα*), and naïve status (*Tcf7*, *Ccr7*, *Cd4*, *Sell*) (Supplementary Figure 7e-g). The integration of stimulation concentrations into the UMAP plot revealed that cells from the 0 μg/ml group were mostly in the naïve cluster and cells from the 1 μg/ml group were mostly in the activated cluster, validating our analysis (Figure 6f). Notably, naïve CD4^+^ T cells exposed to weak stimulation (0.25 μg/ml) were distributed across diverse clusters spanning from naïve to activated states, indicating that reducing the anti-CD3 concentration led to increased heterogeneity in the transcriptomic profile during T cell activation (Figure 6f). Interestingly, the distribution of the two genotypes differed across activation clusters. *Flot2^CD4^* CD4^+^ T cells exhibited a significantly lower frequency in the priming cluster compared to *Flot2^WT^* CD4^+^ T cells (Figure 6g,h). By contrast, *Flot2^CD4^* CD4^+^ T cells demonstrated a higher occupancy in the activated cluster following weak stimulation (0.25 μg/ml) compared to *Flot2^WT^* CD4^+^ T cells, consistent with flow cytometric analysis following *in vitro* stimulation (Figure 6b,h).

Next, using RNA velocity analysis, we examined if Flot2 deficiency affected T cell activation trajectories following *in vitro* stimulation. Our observations revealed nearly overlapping RNA velocity UMAP space between *Flot2^WT^*and *Flot2^CD4^* CD4^+^ T cells, indicating that both groups undergo similar major transcriptional changes during the early stages of activation, regardless of Flot2 expression (Supplementary Figure 7h-j). Although *Flot2^WT^*and *Flot2^CD4^* CD4^+^ T cells displayed similar transcriptomic changes throughout activation, our analysis revealed enhanced RNA splicing of genes associated with cellular proliferation in *Flot2^CD4^*CD4^+^ T cells (Figure 6i,j). Specifically, even in the naive state, *Flot2^CD4^*CD4^+^ T cells demonstrated higher expression levels of spliced *Ppia*, *Cd52*, and *Malat1* (Figure 6j). These genes are known to positively regulate T cell proliferation, activation, as well as to enhance cytotoxic T cell differentiation and cytokine production(45–50). This suggests that while gene expression alterations remain consistent, differences in isoform usage may contribute to variations in T cell activation between *Flot2^WT^* and *Flot2^CD4^* CD4^+^ T cells.

Collectively, these findings suggest that Flot2 deficiency sensitizes the TCR triggering threshold in CD4^+^ T cells, resulting in enhanced TCR signaling and activation, particularly in response to weak TCR stimulation. Flot2 deficiency does not alter the intrinsic activation trajectory of CD4^+^ T cells, but appears to alter the occupancy of a priming state, promoting the progression from the naïve to the fully activated state upon weak stimulation.

### Flot2 controls TCR nanoclustering on the plasma membrane of naïve CD4^+^ T cells

Receptor clustering is pivotal for setting thresholds in various signaling pathways (51, 52). While TCRs form nanoclusters and their clustering is crucial for TCR signaling regulation (7–9, 53–56), the mechanisms governing TCR nanoclustering remain unclear. Based on our findings demonstrating a role of Flot2 in sensitizing TCR signaling initiation, we hypothesized that Flot2 may regulate TCR nanoclustering. Utilizing super-resolution imaging, we examined TCR nanoclusters in steady-state naïve CD4^+^ T cells from *Flot2^WT^*or *Flot2^CD4^* mice, identifying them with CD3ε or TCRβ markers as previously described (9, 57). Notably, we found that naïve *Flot2^CD4^*CD4^+^ T cells exhibited a higher number of CD3ε^+^ TCR nanoclusters in the steady state (Figure 7a-c). Associated with this was a reduction in cluster size, as measured using volumetric space (voxel) analysis (Figure 7d). This increased number of small clusters resulted in a pattern of scattered TCR clusters on the membrane, giving the appearance of greater overall coverage (Figure 7a). Consistent with CD3ε^+^ nanocluster analysis, there was an increase in the number of TCRβ^+^ nanoclusters of smaller sizes (Figure 7e-h). Convex hull geometry analysis further confirmed reduced volume (h.Vol), surface (h.Surface), the largest length (h.Length), and the largest width (h.pBoxW1) of the convex hull of the cluster in *Flot2^CD4^* naïve CD4^+^ T cells compared to *Flot2^WT^* counterparts (Supplementary Figure 8). In summary, these results suggest that Flot2 ablation regulates TCR nanoclustering by promoting the formation of an increased number of small clusters on the plasma membrane of naïve CD4^+^ T cells.

**Figure 7.**
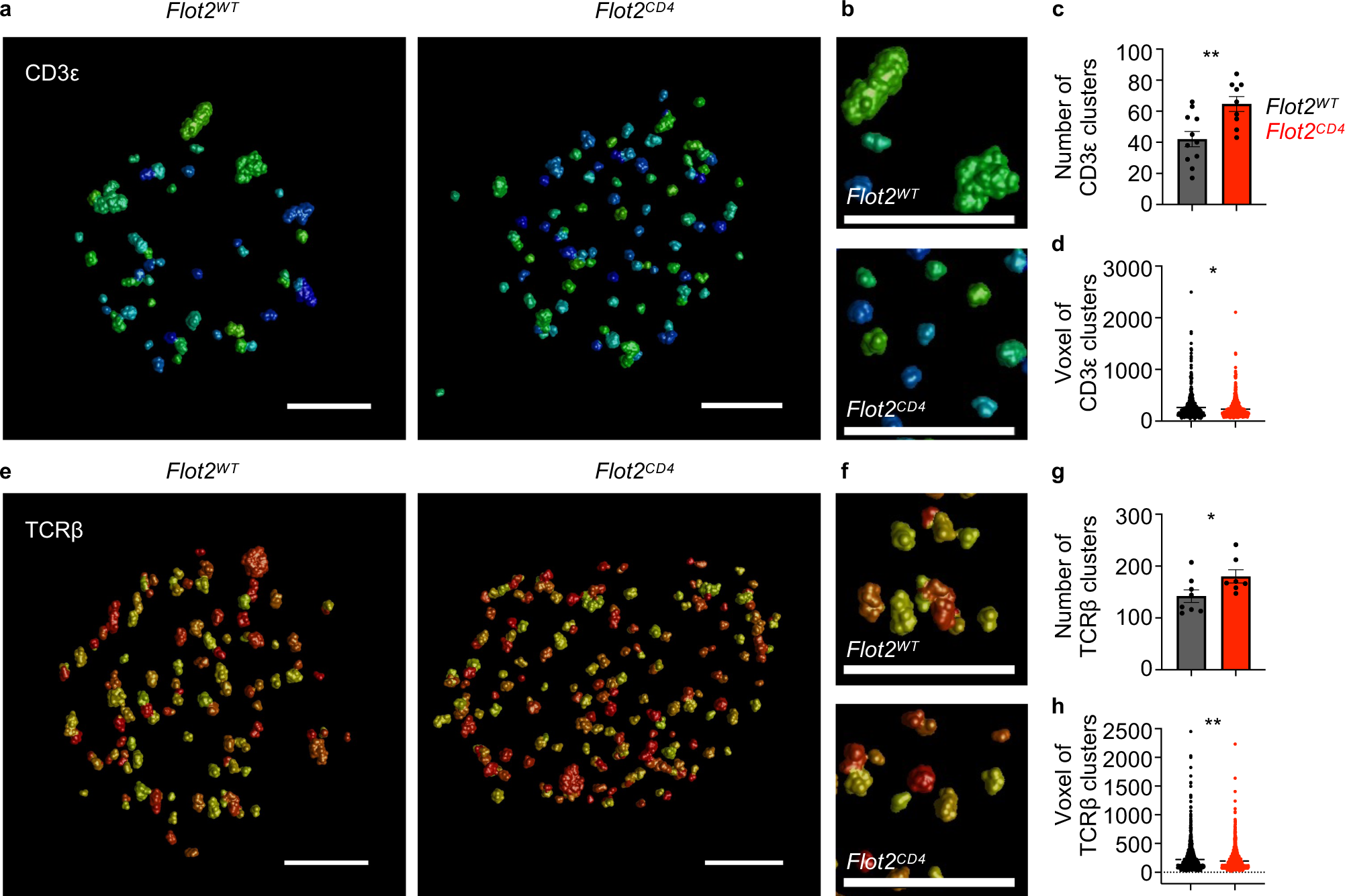
Flot2 ablation increases the number of surface TCR nanoclusters with a smaller size on naïve CD4^+^ T cells. **a**-**h** dSTORM analysis of TCR nanoclustering in *Flot2^WT^* and *Flot2^CD4^* naïve CD4^+^ T cells: Clustering images of CD3ε molecules (**a**, **b**) or TCRβ molecules are shown (**e**, **f**). Number of clusters and voxel of CD3ε (**c**, **d**) or TCRβ (**g**, **h**) molecules after quantification using Huygens cluster analyzer are depicted. Scale bar, 1 μm. Data are representative of four (**a**-**d**) or two (**e**-**h**) independent experiments. Data were analyzed by unpaired t-test (**c**, **d**, **g**, **h**). Error bars denote SEM; *P<0.05; **P<0.01.

## Discussion

T cell immunity relies on the proper initiation of TCR signaling, but how TCR triggering is regulated on the plasma membrane remains unclear. A few prior reports have implicated flotillins in TCR signaling, but conflicting findings have been noted, likely due to contextual factors such as the varying cell types (primary or cell line), stimulation methods, and experimental models that have been used (27, 28, 58). In the present study, we employed complementary *in vivo* and *in vitro* models of primary CD4^+^ and CD8^+^ T cells and included titrated TCR stimulation to comprehensively profile the role of Flot2 in regulating T cell effector functions, T cell differentiation, T cell activation, TCR triggering, and surface TCR distribution. Through this array of approaches, we have identified that Flot2 modulates the TCR triggering threshold and T cell functional responses in CD4^+^ T cells, potentially by orchestrating the spatial organization of surface TCR molecules (Supplementary Figure 9).

An intriguing observation comparing our *in vivo* and *ex vivo* model systems was that *Flot2^CD4^* CD4^+^ T cells displayed an augmented activation profile consistent with the *in vivo* setting, whereas *Flot2^CD4^*CD8^+^ T cells exhibited no noticeable alteration upon *ex vivo* stimulation.

These findings imply that CD8^+^ T cells, unlike CD4^+^ T cells, may depend on additional *in vivo* signals to demonstrate a Flot2-dependent enhanced activation phenotype. The traditional concept of T cell activation encompasses the activation of both CD4^+^ and CD8^+^ T cells via TCR stimulation (signal 1) and costimulation (signal 2)(59, 60). It remains unclear to what extent distinct regulatory mechanisms govern TCR triggering in CD4^+^ versus CD8^+^ T cells, although a few prior reports have identified some cell type-specific differences, including in the impact of IL-2 on TCR signaling threshold, and in the reliance of TCR signaling upon select adaptor proteins (61, 62). Given that Flot2 regulates ganglioside trafficking from the plasma membrane (63), and CD4^+^ and CD8^+^ T cells reportedly have different lipid raft ganglioside composition (64) as well as differential dependence on specific gangliosides for TCR activation (65), it is possible that Flot2 differentially impacts signaling in the two T cell subsets via effects on membrane lipids. It is also possible that CD8^+^ T cells require supplementary Flot2-interacting cues, such as alternative costimulatory signals, integrin engagements, or cytokine effects, during their interactions with antigen-presenting cells. Exploring potential divergent regulatory mechanisms in the initial TCR triggering of CD4^+^ and CD8^+^ T cells presents an interesting avenue for future investigation. Such studies may have important translational value, given that CD4^+^ and CD8^+^ chimeric antigen receptor-expressing T cells used in cancer therapy have distinct functional responses to TCR stimulation (66).

A previous study demonstrated that the strength of stimulation does not inherently determine the activation pathway of T cells; rather, it regulates the speed and synchronicity of cellular activation initiation (44). Considering this, the observation that *Flot2^CD4^*CD4^+^ T cells experienced stronger TCR signaling upon weak stimulation, while overall transcriptional pathways were comparable to WT, suggests that Flot2 deficiency might expedite the transition of cells from a naive state to full activation, rather than altering the T cell activation pathway.

This interpretation is further supported by the reciprocal reduced and increased occupancy of *Flot2^CD4^* CD4^+^ T cells in the priming and activation clusters, respectively, upon weak stimulation. Additionally, the comparable transcriptional activation route suggests that Flot2-mediated regulation of TCR triggering might entail a distinct mechanism, potentially associated with protein spatial organization rather than gene expression regulation. This interpretation is supported by prior reports demonstrating the role of spatial organization of TCR molecules in controlling TCR triggering (7–9), along with our data showing significant differences in the number and size of TCR nanoclusters on naïve CD4^+^ T cells depending on Flot2 expression.

Further investigation is required to elucidate the precise mechanism of how the increased number of smaller TCR nanoclusters sensitizes the activation threshold. Additionally, understanding the impact of Flot2 on the formation of higher-order TCR clusters to create the T cell immunological synapse upon T cell activation will be of great interest.

We observed differential RNA splicing in Flot2-deficient CD4^+^ T cells in the naïve, priming and activated states, in particular, involving genes that regulate cell proliferation, activation, and protein translation (i.e., ribosomal protein-encoding genes). Taken together with our finding of increased levels of activation/maturation markers in lymph node CD4^+^ T cells of naïve *Flot2^CD4^* mice (e.g., Nur77, CD44, T-bet, LFA-1α [Supplementary Figure 3]), this may suggest that increased homeostatic TCR signaling in Flot2-deficient T cells primes alternate isoform usage in genes that support activation and proliferation upon subsequent TCR engagement. In support of the possibility that alternative RNA splicing can regulate TCR signal strength, it was recently reported that the splicing factor SRSF1 regulates T cell activation (67) and that alternative splicing of the adaptor protein MALT1 in response to TCR engagement controls CD4^+^ T cell activation (68).

Immune checkpoint blockade, a cornerstone of anti-cancer therapy, aims to inhibit T cell inhibitory receptors like PD-1 and CTLA-4 (69). However, this approach can lead to side effects such as T cell functional exhaustion and immune-related adverse events due to T cell overactivation (70–72). Our findings reveal that Flot2 deficiency enhances T cell responsiveness to low TCR stimulation, resulting in improved effector responses and tumor control in an *in vivo* tumor model, while mitigating T cell functional exhaustion. Importantly, neither global nor T cell-specific Flot2 knockout mice exhibited the spontaneous autoimmune phenotypes seen in PD-1 or CTLA-4 knockout mice (73–78), suggesting that therapeutic Flot2 targeting may be less subject to deleterious immunological side effects. Given its unique mechanism of regulating TCR nanocluster formation, Flot2 deficiency may hold promise for providing synergy in combination therapies with existing anticancer T cell approaches.

In conclusion, the present study demonstrates that Flot2 deletion can boost T cell antigen sensitivity as well as T cell effector functionality, potentially through regulating surface TCR clustering. These findings suggest that targeting Flot2 either *in vivo* or through engineering of T cells for adoptive cell therapy may hold promise for enhancing T cell reactivity in diseases with weak antigenicity, including cancer and chronic infections. We also posit that our findings suggest future avenues for investigating membrane-level mechanisms of receptor nanoclustering and also for understanding differences in TCR signal transduction between CD4^+^ and CD8^+^ T cells, and potentially for how these differences may be leveraged therapeutically.

## Methods

### Sex as a biological variable

*In vivo* tumor studies were conducted using mice of both sexes, with sex-matched controls. Similar results were observed in both sexes, as described in the Results.

### Mice

*Flot2^flox/flox^* mice were generated by insertion of loxP sites flanking coiled-coil domain(14, 79) of the *Flot2* locus using standard cloning and homologous recombination methods involving electroporation of the targeting vector into mouse embryonic stem (ES) cells, followed by ES screening, blastocyst injection of appropriately targeted ES cells, and then breeding of male chimeras with wild-type C57BL/6 females. Subsequent global *Flot2* deletion was achieved by crossbreeding with CMV-Cre mice (Jackson Laboratory #006054), and T cell-specific deletion was achieved by crossbreeding with *CD4^cre^* transgenic mice (Jackson Laboratory #022071). All experiments used sex-matched controls of age ∼6-12 weeks.

### *In vitro* stimulation of naïve T cells

Naïve CD4^+^ or CD8^+^ T cells were isolated from mouse spleen and lymph nodes by using a mouse naïve CD4^+^ or CD8^+^ T cell isolation kit (STEMCELL Technologies). Purified naïve CD4^+^ or CD8^+^ T cells were labeled with CellTrace Violet (Invitrogen) and cultured in RPMI-1640 (Gibco) complete medium supplemented with 10% fetal bovine serum, 57.2 μM β-mercaptoethanol, and 1X antibiotic-antimycotic (Gibco) at a density of 1 x 10^5^ cells per well in a 96 well plate. Naïve CD4^+^ or CD8^+^ T cells were stimulated with various concentrations of plate-bound anti-CD3 and a fixed concentration of soluble anti-CD28 (1 μg/ml) antibodies for different time points as indicated in the figure legends.

### *In vitro* polarization of T helper cells

Naïve CD4^+^ T cells were purified from mouse spleen and lymph nodes by using a mouse naïve CD4^+^ T cell isolation kit (STEMCELL Technologies) and polarized following the manufacturer’s protocol for Immunocult Mouse Th1 or Th2 differentiation supplements (STEMCELL Technologies). For Th17 polarization, naïve CD4^+^ T cells were cultured with 5 ng/ml TGF-β and 20 ng/ml IL-6, while Treg polarization was induced by culturing cells with 5 ng/ml rhIL-2 and 5 ng/ml TGF-β, both for 3.5 days. All T helper cell polarization involved cell stimulation using the indicated concentration of plate-bound anti-CD3 and 0.5 μg/ml soluble anti-CD28. Polarization efficiency was assessed by quantifying cytokine production and lineage marker expression via flow cytometry following PMA/ionomycin stimulation with protein transport inhibitors (Invitrogen) for 4 hours at 37°C.

### Single cell RNA sequencing and data processing

Single cell suspensions of stimulated *Flot2^WT^* or *Flot2^CD4^* naïve CD4^+^ T cells were washed and loaded and processed for 10X Genomics scRNA-Seq analysis. Fastq files were processed using cellranger’s “multi” functionality for hashtag oligos using information of the two separate sequencing outputs, cellranger’s mm10-3.0.0 reference, and associated oligo hashtag information. Data were analyzed using Seurat v5.0.1 (80) in R4.3.1 as previously (81). In short, the scRNA-Seq dataset from two sample lanes was merged with Hashtag information (three antibodies per sample lane). Hashtag information was demultiplexed and classified using the “HTODemux” function, and subset only on cells that were assigned singlet classification, to generate six separate samples in total. Samples were then filtered for homogeneity, using the following parameters: nFeature_RNA (500-4,250), nCount_RNA (100-20,000), percent mitochondria (0.0075 – 0.0800), percent cycling (0.0075 – 0.0800). Count matrices were normalized and scaled for number of RNA features, proportion cycling and proportion mitochondrial content. Subsequently, data dimensions were reduced by PCA and by UMAP (using the first 30 PCs). Clustering was performed using the “FindClusters” function using the Louvain algorithm (k.param = 50, resolution = 0.5). Genes identifying cluster membership were generated by the “FindAllMarkers” function.

Spliced and unspliced count matrices were generated with the “velocyto run” function using the same reference genes as cellranger (see above) and bam files generated from cellranger(82). RNA velocity results were displayed using velocyto.R v0.6 (82).

### Super-resolution imaging and analysis of TCR nanoclusters

To visualize TCR distribution in the plasma membrane, staining was conducted as previously described with a modification (57). Briefly, 5 × 10^5^ *Flot2^WT^* or *Flot2^CD4^* naïve CD4^+^ T cells were fixed with 4% PFA, followed by surface staining with 5 μg/ml anti-mCD3ε for 4 hours at 4°C. Subsequently, the cells were stained with 2 μg/ml Alexa 647-conjugated goat anti-hamster IgG for 2 hours at 4°C after washing with PBS ten times. In cases where Alexa 647-conjugated anti-mTCRβ antibody was used, secondary staining was omitted. Stained cells were resuspended in 40 μl PBS and transferred to the poly-D-lysine coated dish, followed by overnight incubation at 37°C for cell attachment to the dish bottom. A mercaptoethylamine-based STORM cocktail was used was for super resolution imaging through localization microscopy. Three solutions were made consisting of: Solution A (0.8mL) – 30mM Tris/Cl pH 8.5 containing 1 mM EDTA and 6.25 μM glucose oxidate + 2.5 μM catalase, Solution B (0.1mL) – 250 mM cysteamine-HCL in water, and Solution C (0.1mL) – 250 mM glucose in water. Solution A was gently mixed with Solution B, this combined mixture was gently added to Solution C. This ABC mixture was immediately pulled into a gas-tight glass syringe with a PEEK needle minimizing any air bubbles. The PBS solution from the fixed/labeled cells attached to the poly-D-lysine coated dish was removed, and a 25 mm square coverslip was placed over the 14 mm microwell and then the PEEK needle from the glass syringe was used to inject the ABC solution across the sample to deplete dissolved oxygen from the sample chamber. Ten thousand images were captured in burst mode on an Andor Dragonfly 505 imaging system using its 3D astigmatic lens for 3D super-resolution imaging. A 637 nm laser at 100% with the PD4 power density setting was used to excite Alexa647 through a Nikon CFI Aprochromat TIRF 60X Oil Immersion lens and the corresponding fluorescence emission was captured through a 660-738 nm emission filter and Andor iXon EMCCD camera with an exposure of 10msec. This image series was then taken into Huygens Localizer (v22.10, Scientific Volume Imaging, Hilversum, Netherlands) for generation of a 3D localization table using the Weighted Least-squares fit method with a 3D Z-position calibration PSF and drift correction applied. The 3D localization table was then opened with Huygen’s Cluster Analyzer 22.10 or 23.10, where the FOCAL algorithm (nearestNeighborsCross 7, threshold 20, and minimal voxel count 9) or the DBSCAN algorithm (a minimal neighbors 6 and minimal cluster size 1), were used to identify CD3ε or TCRβ clusters, respectively.

### Listeria monocytogenes infection

Mice were infected with *L. monocytogenes* via retro-orbital intravenous injection, receiving a dosage of 5000 colony-forming units (CFU) per mouse. Daily monitoring of weight loss post-infection was conducted and subsequently analyzed. Spleens were excised and mechanically dissociated to obtain total splenocytes. Splenocytes were counted and then stimulated with PMA/ionomycin in the presence of protein transport inhibitors (Invitrogen) for 4 hours at 37 °C. Subsequently, FACS analysis was performed to measure T cell effector cytokine production and Ki67 expression.

### Tumor models

5 x 10^5^ B16F10 or 5 x 10^5^ MC38 cells were injected intradermally into *Flot2^+/+^* and *Flot2^−/−^* mice, and subcutaneously into *Flot2^WT^* and *Flot2^CD4^* mice. From day 7 or day 8, tumor size was monitored every 2 days. Tumor size was calculated as (length × width^2^)/2. Tumor-bearing mice were monitored and analyzed until day 20. The experimental endpoint was determined based on tumor size (with a limit of 2000 mm^3^), severe tumor ulcerations, or other health issues that conflicted with the approved animal study protocol by the Animal Care and Use Committee of the NIEHS. Tumor and draining lymph nodes were excised and mechanically dissociated, and T cells in each tissue were analyzed by FACS.

### Mixed bone-marrow chimera

Mixed bone-marrow chimera experiments were performed as previously described(34). Briefly, bone-marrow from either WT or *Flot2^−/−^* mice were mixed with bone-marrow from *TCRα^−/−^* mice at a 1:5 ratio and transferred into *TCRα^−/−^* recipient mice that were lethally irradiated at Rad 1,100. After 10 weeks of reconstitution, 2 x 10^5^ B16F10 tumor cells were intradermally injected, and tumor growth and TILs were analyzed.

### Quantitative PCR

Total RNA was obtained using RNeasy kits (Qiagen) following manufacturer’s instructions. cDNA was synthesized using the iScript cDNA synthesis kit (Bio-Rad). cDNA was quantified with TaqMan Universal PCR Master Mix (Invitrogen) and predesigned TaqMan primers (Assay ID: Mm00514962_g1, Mm01241315_g1). The delta-delta Ct method was utilized for analyzing the fold change in gene expression, which was normalized using appropriate reference genes. QuantStudio Software (ThermoFisher Scientific) was used for data analysis.

### Reagents

For the flow cytometric analysis, following reagents were used: CellTrace™ Violet Cell Proliferation Kit (Invitrogen, C34557A), 7AAD (Sigma, A9400-1MG), anti-CD45.2 (BD Bioscience, 612778), anti-CD45 (Invitrogen, 48-0451-82), anti-TCRβ (Biolegend, 109246; BD Biosciences, 553170), anti-CD4 (Biolegend, 100549), anti-CD8α (Biolegend, 100741), anti-CD62L (eBioscience, 12-0621-81; Cytek, 60-0621-U025; Biolegend, 104436), anti-CD44 (Biolegend, 103047 or 740215), anti-CD69 (Biolegend, 104512), anti-CD25 (Biolegend, 102017; eBioscience, 47-0251-82), anti-Nur77 (Invitrogen, 12-5965-82), anti-Ki67 (Biolegend, 652413, 652403, or 652411), anti-LFA-1 (Biolegend, 141012), anti-T-bet (Biolegend, 644832; eBioscience, 53-5825-82), anti-TCF1 (Cell Signaling, 6709S or 90511S), anti-TOX (Cell Signaling, 44682S), anti-CXCR5 (Biolegend, 145522), anti-TIM3 (Biolegend, 119723), anti-PD1 (BD Biosciences, 744544; Biolegend, 135220), anti-Foxp3 (eBioscience, 12-4771-82; Biolegend, 126406), anti-TNFα (BioLegend, 506308), anti-IFNγ (Biolegend, 505826), anti-IL-2 (BD Biosciences, 557725), anti-IL-17 (BD Biosciences, 559502), anti-IL-10 (Biolegend, 505016), anti-GraB (Biolegend, 372204). For western blots, the following reagents were used: NuPAGE LDS sample buffer (Invitrogen, NP0007), 4–12% Criterion™ XT Bis-Tris Protein Gel (Bio-Rad, 3450125), NuPAGE™ MOPS SDS Running Buffer (Invitrogen, NP0001), Precision Plus Protein™ Kaleidoscope™ Prestained Protein Standards (Bio-Rad, 1610375), iBlot™ 2 Transfer Stacks (Invitrogen, IB24001), Phospho-Lck (Tyr505) Antibody (Cell Signaling, 2751), Phospho-Zap-70 (Tyr319)/Syk (Tyr352) (Cell Signaling, 2717), Lck Antibody (Cell Signaling, 2752), Anti-ZAP-70 Kinase (BD Biosciences, 610239), Goat anti-Rabbit IgG (H+L) Poly-HRP Secondary Antibody HRP (Invitrogen, 32260), Anti-mouse IgG HRP-linked Antibody (Cell Signaling, 7076), Anti-β-Actin−Peroxidase antibody (Sigma-Aldrich, A3854), Clarity Western ECL Substrate (Bio-Rad, 1705061), Restore™ Western Blot Stripping Buffer (Thermo Scientific, 21059). For scRNA-seq sample labeling, following antibodies were used: TotalSeq™-A0301 anti-mouse Hashtag 1 Antibody (Biolegend, 155801), TotalSeq™-A0302 anti-mouse Hashtag 2 Antibody (Biolegend, 155803), and TotalSeq™-A0303 anti-mouse Hashtag 3 Antibody (Biolegend, 155805).

### Statistical analysis

Statistical analyses were conducted using GraphPad Prism (GraphPad Software, Inc.), with significance levels as specified in the figure legends.

### Study approval

All experiments were performed in accordance with the Animal Welfare Act and the US Public Health Service Policy on Humane Care and Use of Laboratory Animals after review by the Animal Care and Use Committee of the National Institute of Environmental Health Sciences (NIEHS).

## Data Availability

Sc-RNA-Seq expression data will be made available on GEO.

## Author contributions

S.M. conceptualized the study, conducted experiments, analyzed the data, and drafted the initial manuscript. F.Z. also conceived the study, conducted experiments, and participated in data analysis. M.U. conducted western blot experiments, while C.T. aided in data acquisition and analysis of super-resolution imaging. P.K. analyzed scRNA-seq data and offered intellectual input on experimental design and analysis in general. M.F. supervised the project, contributed to experimental design and analysis, and provided overall guidance. All authors contributed to manuscript drafting.

## Acknowledgments

These studies were supported by the Intramural Research Program of the National Institute of Environmental Health Sciences, NIH (Z01 ES102005). The authors are grateful to Ms. Ligon Perrow for assistance with animal husbandry as well as to the staff of the NIEHS Flow Cytometry Core Facility and the NIEHS Fluorescence Microscopy and Imaging Center.

## References

1. McKeithan TW. Kinetic proofreading in T-cell receptor signal transduction. Proceedings of the National Academy of Sciences. 1995;92(11):5042–6.

2. Lo WL, Shah NH, Rubin SA, Zhang W, Horkova V, Fallahee IR, et al. Slow phosphorylation of a tyrosine residue in LAT optimizes T cell ligand discrimination. Nat Immunol. 2019;20(11):1481–93.

3. Pettmann J, Huhn A, Abu Shah E, Kutuzov MA, Wilson DB, Dustin ML, et al. The discriminatory power of the T cell receptor. eLife. 2021;10:e67092.

4. Tischer DK, and Weiner OD. Light-based tuning of ligand half-life supports kinetic proofreading model of T cell signaling. eLife. 2019;8:e42498.

5. Voisinne G, Locard-Paulet M, Froment C, Maturin E, Menoita MG, Girard L, et al. Kinetic proofreading through the multi-step activation of the ZAP70 kinase underlies early T cell ligand discrimination. Nature Immunology. 2022;23(9):1355–64.

6. Yousefi OS, Günther M, Hörner M, Chalupsky J, Wess M, Brandl SM, et al. Optogenetic control shows that kinetic proofreading regulates the activity of the T cell receptor. eLife. 2019;8:e42475.

7. Taylor MJ, Husain K, Gartner ZJ, Mayor S, and Vale RD. A DNA-Based T Cell Receptor Reveals a Role for Receptor Clustering in Ligand Discrimination. Cell. 2017;169(1):108–19.e20.

8. Cai H, Muller J, Depoil D, Mayya V, Sheetz MP, Dustin ML, and Wind SJ. Full control of ligand positioning reveals spatial thresholds for T cell receptor triggering. Nature Nanotechnology. 2018;13(7):610–7.

9. Pageon SV, Tabarin T, Yamamoto Y, Ma Y, Nicovich PR, Bridgeman JS, et al. Functional role of T-cell receptor nanoclusters in signal initiation and antigen discrimination. Proceedings of the National Academy of Sciences. 2016;113(37):E5454–E63.

10. Bickel PE, Scherer PE, Schnitzer JE, Oh P, Lisanti MP, and Lodish HF. Flotillin and Epidermal Surface Antigen Define a New Family of Caveolae-associated Integral Membrane Proteins*. Journal of Biological Chemistry. 1997;272(21):13793–802.

11. Schulte T, Paschke KA, Laessing U, Lottspeich F, and Stuermer CAO. Reggie-1 and reggie-2, two cell surface proteins expressed by retinal ganglion cells during axon regeneration. Development. 1997;124(2):577–87.

12. Volonté D, Galbiati F, Li S, Nishiyama K, Okamoto T, and Lisanti MP. Flotillins/Cavatellins Are Differentially Expressed in Cells and Tissues and Form a Hetero-oligomeric Complex with Caveolins in Vivo. Journal of Biological Chemistry. 1999;274(18):12702–9.

13. Rivera-Milla E, Stuermer CAO, and Málaga-Trillo E. Ancient origin of reggie (flotillin), reggie-like, and other lipid-raft proteins: convergent evolution of the SPFH domain. Cellular and Molecular Life Sciences CMLS. 2006;63(3):343–57.

14. Gonzalo, Hoegg M, Munderloh C, Schrock Y, Malaga-Trillo E, Rivera-Milla E, and Claudia. Reggie/flotillin proteins are organized into stable tetramers in membrane microdomains. Biochemical Journal. 2007;403(2):313–22.

15. Munderloh C, Solis GP, Bodrikov V, Jaeger FA, Wiechers M, Málaga-Trillo E, and Stuermer CAO. Reggies/Flotillins Regulate Retinal Axon Regeneration in the Zebrafish Optic Nerve and Differentiation of Hippocampal and N2a Neurons. The Journal of Neuroscience. 2009;29(20):6607–15.

16. Baumgart T, Hammond AT, Sengupta P, Hess ST, Holowka DA, Baird BA, and Webb WW. Large-scale fluid/fluid phase separation of proteins and lipids in giant plasma membrane vesicles. Proceedings of the National Academy of Sciences. 2007;104(9):3165–70.

17. Diaz-Rohrer BB, Levental KR, Simons K, and Levental I. Membrane raft association is a determinant of plasma membrane localization. Proceedings of the National Academy of Sciences. 2014;111(23):8500–5.

18. Varshney P, Yadav V, and Saini N. Lipid rafts in immune signalling: current progress and future perspective. Immunology. 2016;149(1):13–24.

19. Kulkarni R, Wiemer EAC, and Chang W. Role of Lipid Rafts in Pathogen-Host Interaction - A Mini Review. Frontiers in Immunology. 2022;12.

20. Wong SW, Kwon M-J, Choi AMK, Kim H-P, Nakahira K, and Hwang DH. Fatty Acids Modulate Toll-like Receptor 4 Activation through Regulation of Receptor Dimerization and Recruitment into Lipid Rafts in a Reactive Oxygen Species-dependent Manner. Journal of Biological Chemistry. 2009;284(40):27384–92.

21. Lafont F, and Simons K. Raft-partitioning of the ubiquitin ligases Cbl and Nedd4 upon IgE-triggered cell signaling. Proceedings of the National Academy of Sciences. 2001;98(6):3180–4.

22. Cheng PC, Dykstra ML, Mitchell RN, and Pierce SK. A Role for Lipid Rafts in B Cell Antigen Receptor Signaling and Antigen Targeting. The Journal of Experimental Medicine. 1999;190(11):1549–60.

23. Cherukuri A, Shoham T, Sohn HW, Levy S, Brooks S, Carter R, and Pierce SK. The Tetraspanin CD81 Is Necessary for Partitioning of Coligated CD19/CD21-B Cell Antigen Receptor Complexes into Signaling-Active Lipid Rafts. The Journal of Immunology. 2004;172(1):370–80.

24. Drevot P. TCR signal initiation machinery is pre-assembled and activated in a subset of membrane rafts. The EMBO Journal. 2002;21(8):1899–908.

25. Tavano R, Contento RL, Baranda SJ, Soligo M, Tuosto L, Manes S, and Viola A. CD28 interaction with filamin-A controls lipid raft accumulation at the T-cell immunological synapse. Nature Cell Biology. 2006;8(11):1270–6.

26. Tavano R, Gri G, Molon B, Marinari B, Rudd CE, Tuosto L, and Viola A. CD28 and Lipid Rafts Coordinate Recruitment of Lck to the Immunological Synapse of Human T Lymphocytes. The Journal of Immunology. 2004;173(9):5392–7.

27. Compeer EB, Kraus F, Ecker M, Redpath G, Amiezer M, Rother N, et al. A mobile endocytic network connects clathrin-independent receptor endocytosis to recycling and promotes T cell activation. Nature Communications. 2018;9(1).

28. Yingqiu Li CZ. In: University SYS ed. China; 2023.

29. Schwenk F, Baron U, and Rajewsky K. A *cre*-transgenic mouse strain for the ubiquitous deletion of *loxP*-flanked gene segments including deletion in germ cells. Nucleic Acids Research. 1995;23(24):5080–1.

30. Overwijk WW, and Restifo NP. B16 as a Mouse Model for Human Melanoma. Current Protocols in Immunology. 2000;39(1).

31. Gordon SR, Maute RL, Dulken BW, Hutter G, George BM, McCracken MN, et al. PD-1 expression by tumour-associated macrophages inhibits phagocytosis and tumour immunity. Nature. 2017;545(7655):495-9.

32. Olson B, Li Y, Lin Y, Liu ET, and Patnaik A. Mouse Models for Cancer Immunotherapy Research. Cancer Discovery. 2018;8(11):1358–65.

33. Dranoff G. Experimental mouse tumour models: what can be learnt about human cancer immunology? Nature Reviews Immunology. 2012;12(1):61–6.

34. Karmaus PWF, Chen X, Lim SA, Herrada AA, Nguyen TM, Xu B, et al. Metabolic heterogeneity underlies reciprocal fates of T(H)17 cell stemness and plasticity. Nature. 2019;565(7737):101–5.

35. Maggioli MF, Palmer MV, Thacker TC, Vordermeier HM, McGill JL, Whelan AO, et al. Increased TNF-α/IFN-γ/IL-2 and Decreased TNF-α/IFN-γ Production by Central Memory T Cells Are Associated with Protective Responses against Bovine Tuberculosis Following BCG Vaccination. Frontiers in Immunology. 2016;7.

36. Westerhof LM, McGuire K, Maclellan L, Flynn A, Gray JI, Thomas M, et al. Multifunctional cytokine production reveals functional superiority of memory CD4 T cells. European Journal of Immunology. 2019;49(11):2019–29.

37. Zhu J, Yamane H, and Paul WE. Differentiation of Effector CD4 T Cell Populations. Annual Review of Immunology. 2010;28(1):445–89.

38. Saravia J, Chapman NM, and Chi H. Helper T cell differentiation. Cellular & Molecular Immunology. 2019;16(7):634–43.

39. Zhu X, and Zhu J. CD4 T Helper Cell Subsets and Related Human Immunological Disorders. International Journal of Molecular Sciences. 2020;21(21):8011.

40. Bartleson JM, Viehmann Milam AA, Donermeyer DL, Horvath S, Xia Y, Egawa T, and Allen PM. Strength of tonic T cell receptor signaling instructs T follicular helper cell–fate decisions. Nature Immunology. 2020;21(11):1384–96.

41. Bhattacharyya ND, and Feng CG. Regulation of T Helper Cell Fate by TCR Signal Strength. Frontiers in Immunology. 2020;11.

42. Snook JP, Kim C, and Williams MA. TCR signal strength controls the differentiation of CD4 ^+^ effector and memory T cells. Science Immunology. 2018;3(25):eaas9103.

43. Li MO, and Rudensky AY. T cell receptor signalling in the control of regulatory T cell differentiation and function. Nature Reviews Immunology. 2016;16(4):220–33.

44. Richard AC, Lun ATL, Lau WWY, Göttgens B, Marioni JC, and Griffiths GM. T cell cytolytic capacity is independent of initial stimulation strength. Nature Immunology. 2018;19(8):849–58.

45. Nigro P, Pompilio G, and Capogrossi MC. Cyclophilin A: a key player for human disease. Cell Death & Disease. 2013;4(10):e888-e.

46. Han JM, and Jung HJ. Cyclophilin A/CD147 Interaction: A Promising Target for Anticancer Therapy. International Journal of Molecular Sciences. 2022;23(16):9341.

47. Ramouz A, Nikbakhsh R, Khajeh E, Sadeghi M, Daniel V, Schmitzler P, et al. Preoperative CD52 Level Predicts Graft Survival following Kidney Transplantation. BioMed Research International. 2022;2022:1–7.

48. Zhao Y, Su H, Shen X, Du J, Zhang X, and Zhao Y. The immunological function of CD52 and its targeting in organ transplantation. Inflammation Research. 2017;66(7):571–8.

49. Wheeler BD, Gagnon JD, Zhu WS, Muñoz-Sandoval P, Wong SK, Simeonov DR, et al.: eLife Sciences Publications, Ltd; 2023.

50. Tripathi V, Shen Z, Chakraborty A, Giri S, Freier SM, Wu X, et al. Long Noncoding RNA MALAT1 Controls Cell Cycle Progression by Regulating the Expression of Oncogenic Transcription Factor B-MYB. PLoS Genetics. 2013;9(3):e1003368.

51. Li M, and Yu Y. Innate immune receptor clustering and its role in immune regulation. Journal of Cell Science. 2021;134(4):jcs249318.

52. Bray D, Levin MD, and Morton-Firth CJ. Receptor clustering as a cellular mechanism to control sensitivity. Nature. 1998;393(6680):85–8.

53. Dong R, Aksel T, Chan W, Germain RN, Vale RD, and Douglas SM. DNA origami patterning of synthetic T cell receptors reveals spatial control of the sensitivity and kinetics of signal activation. Proceedings of the National Academy of Sciences. 2021;118(40):e2109057118.

54. Dushek O, and Van Der Merwe PA. An induced rebinding model of antigen discrimination. Trends in Immunology. 2014;35(4):153–8.

55. Martínez-Martín N, Risueño RM, Morreale A, Zaldívar I, Fernández-Arenas E, Herranz F, et al. Cooperativity Between T Cell Receptor Complexes Revealed by Conformational Mutants of CD3ɛ. Science Signaling. 2009;2(83):ra43-ra.

56. Schamel WWA, and Alarcón B. Organization of the resting TCR in nanoscale oligomers. Immunological Reviews. 2013;251(1):13–20.

57. Yang W, Bai Y, Xiong Y, Zhang J, Chen S, Zheng X, et al. Potentiating the antitumour response of CD8+ T cells by modulating cholesterol metabolism. Nature. 2016;531(7596):651–5.

58. Ficht X, Ruef N, Stolp B, Samson GPB, Moalli F, Page N, et al. In Vivo Function of the Lipid Raft Protein Flotillin-1 during CD8+ T Cell–Mediated Host Surveillance. The Journal of Immunology. 2019;203(9):2377–87.

59. Smith-Garvin JE, Koretzky GA, and Jordan MS. T Cell Activation. Annual Review of Immunology. 2009;27(1):591–619.

60. Tai Y, Wang Q, Korner H, Zhang L, and Wei W. Molecular Mechanisms of T Cells Activation by Dendritic Cells in Autoimmune Diseases. Frontiers in Pharmacology. 2018;9.

61. Au-Yeung BB, Smith GA, Mueller JL, Heyn CS, Jaszczak RG, Weiss A, and Zikherman J. IL-2 Modulates the TCR Signaling Threshold for CD8 but Not CD4 T Cell Proliferation on a Single-Cell Level. The Journal of Immunology. 2017;198(6):2445–56.

62. Parzmair GP, Gereke M, Haberkorn O, Annemann M, Podlasly L, Kliche S, et al. ADAP plays a pivotal role in CD4+ T cell activation but is only marginally involved in CD8+ T cell activation, differentiation, and immunity to pathogens. Journal of Leukocyte Biology. 2017;101(2):407–19.

63. Saslowsky DE, Cho JA, Chinnapen H, Massol RH, Chinnapen DJF, Wagner JS, et al. Intoxication of zebrafish and mammalian cells by cholera toxin depends on the flotillin/reggie proteins but not Derlin-1 or -2. Journal of Clinical Investigation. 2010;120(12):4399–409.

64. de Mello Coelho V, Nguyen D, Giri B, Bunbury A, Schaffer E, and Taub DD. Quantitative differences in lipid raft components between murine CD4+ and CD8+ T cells. BMC Immunol. 2004;5:2.

65. Nagafuku M, Okuyama K, Onimaru Y, Suzuki A, Odagiri Y, Yamashita T, et al. CD4 and CD8 T cells require different membrane gangliosides for activation. Proceedings of the National Academy of Sciences. 2012;109(6):E336–E42.

66. Yang Y, Kohler ME, Chien CD, Sauter CT, Jacoby E, Yan C, et al. TCR engagement negatively affects CD8 but not CD4 CAR T cell expansion and leukemic clearance. Science Translational Medicine. 2017;9(417):eaag1209.

67. Katsuyama T, Li H, Comte D, Tsokos GC, and Moulton VR. Splicing factor SRSF1 controls T cell hyperactivity and systemic autoimmunity. Journal of Clinical Investigation. 2019;129(12):5411–23.

68. Meininger I, Griesbach RA, Hu D, Gehring T, Seeholzer T, Bertossi A, et al. Alternative splicing of MALT1 controls signalling and activation of CD4+ T cells. Nature Communications. 2016;7(1):11292.

69. Korman AJ, Garrett-Thomson SC, and Lonberg N. The foundations of immune checkpoint blockade and the ipilimumab approval decennial. Nature Reviews Drug Discovery. 2022;21(7):509–28.

70. Ramos-Casals M, Brahmer JR, Callahan MK, Flores-Chávez A, Keegan N, Khamashta MA, et al. Immune-related adverse events of checkpoint inhibitors. Nature Reviews Disease Primers. 2020;6(1).

71. Tison A, Garaud S, Chiche L, Cornec D, and Kostine M. Immune-checkpoint inhibitor use in patients with cancer and pre-existing autoimmune diseases. Nature Reviews Rheumatology. 2022;18(11):641–56.

72. Yin Q, Wu L, Han L, Zheng X, Tong R, Li L, et al. Immune-related adverse events of immune checkpoint inhibitors: a review. Frontiers in Immunology. 2023;14.

73. Nishimura H, Nose M, Hiai H, Minato N, and Honjo T. Development of Lupus-like Autoimmune Diseases by Disruption of the PD-1 Gene Encoding an ITIM Motif-Carrying Immunoreceptor. Immunity. 1999;11(2):141–51.

74. Nishimura H, Okazaki T, Tanaka Y, Nakatani K, Hara M, Matsumori A, et al. Autoimmune Dilated Cardiomyopathy in PD-1 Receptor-Deficient Mice. Science. 2001;291(5502):319–22.

75. Okazaki T, Tanaka Y, Nishio R, Mitsuiye T, Mizoguchi A, Wang J, et al. Autoantibodies against cardiac troponin I are responsible for dilated cardiomyopathy in PD-1-deficient mice. Nature Medicine. 2003;9(12):1477–83.

76. Chambers CA, Cado D, Truong T, and Allison JP. Thymocyte development is normal in CTLA-4-deficient mice. Proceedings of the National Academy of Sciences. 1997;94(17):9296–301.

77. Tivol EA, Borriello F, Schweitzer AN, Lynch WP, Bluestone JA, and Sharpe AH. Loss of CTLA-4 leads to massive lymphoproliferation and fatal multiorgan tissue destruction, revealing a critical negative regulatory role of CTLA-4. Immunity. 1995;3(5):541–7.

78. Waterhouse P, Penninger JM, Timms E, Wakeham A, Shahinian A, Lee KP, et al. Lymphoproliferative Disorders with Early Lethality in Mice Deficient in Ctla-4. Science. 1995;270(5238):985–8.

79. Gauthier-Rouvière C, Bodin S, Comunale F, and Planchon D. Flotillin membrane domains in cancer. Cancer and Metastasis Reviews. 2020;39(2):361–74.

80. Hao Y, Stuart T, Kowalski MH, Choudhary S, Hoffman P, Hartman A, et al. Dictionary learning for integrative, multimodal and scalable single-cell analysis. Nature Biotechnology. 2024;42(2):293–304.

81. Izumi G, Nakano H, Nakano K, Whitehead GS, Grimm SA, Fessler MB, et al. CD11b+ lung dendritic cells at different stages of maturation induce Th17 or Th2 differentiation. Nature Communications. 2021;12(1).

82. La Manno G, Soldatov R, Zeisel A, Braun E, Hochgerner H, Petukhov V, et al. RNA velocity of single cells. Nature. 2018;560(7719):494–8.

