## Supplemental Material for "Flotillin-2 dampens T cell antigen-sensitivity and functionality"

### SUPPLEMENTAL MATERIALS

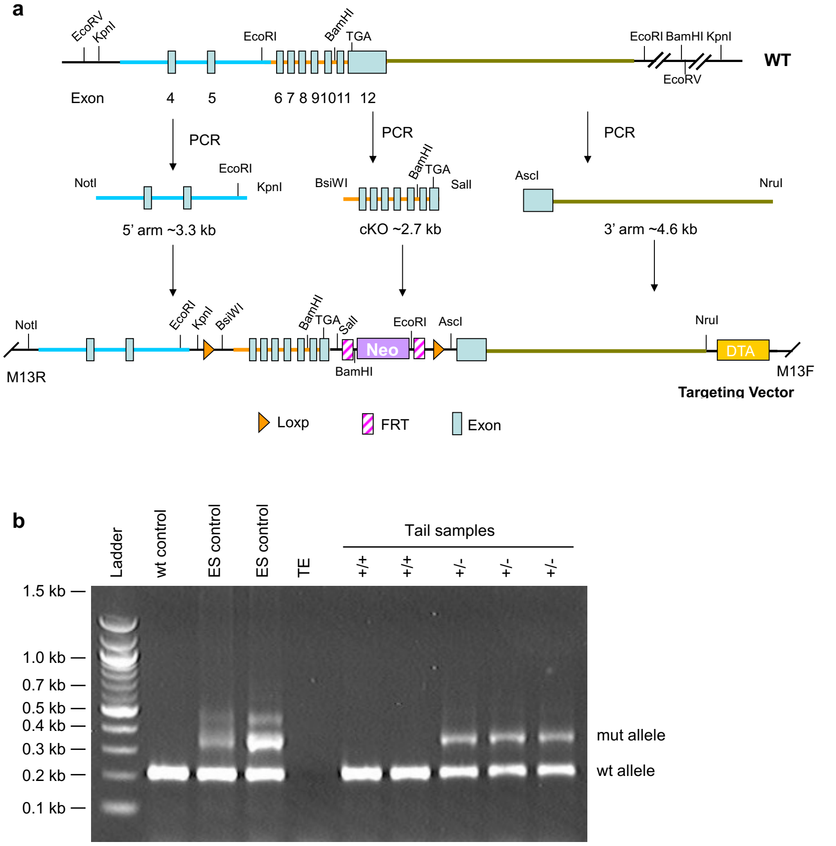


**Supplementary Figure 1**. **Flotillin knockout mouse targeting vector strategy and confirmation.** **a** The vector targeting strategy for flanking the murine *Flot2* locus with loxP sites is shown, along with restriction sites. **b** Pups from chimera breeding were screened by PCR of tail samples using the following primers: 5'-ATCACTGTCTGTCTGTGAGGAGTGG-3' and 5'-AGGGCAAGAGCGTGTGGGTTGTGG-3' followed by gel electrophoresis as shown.

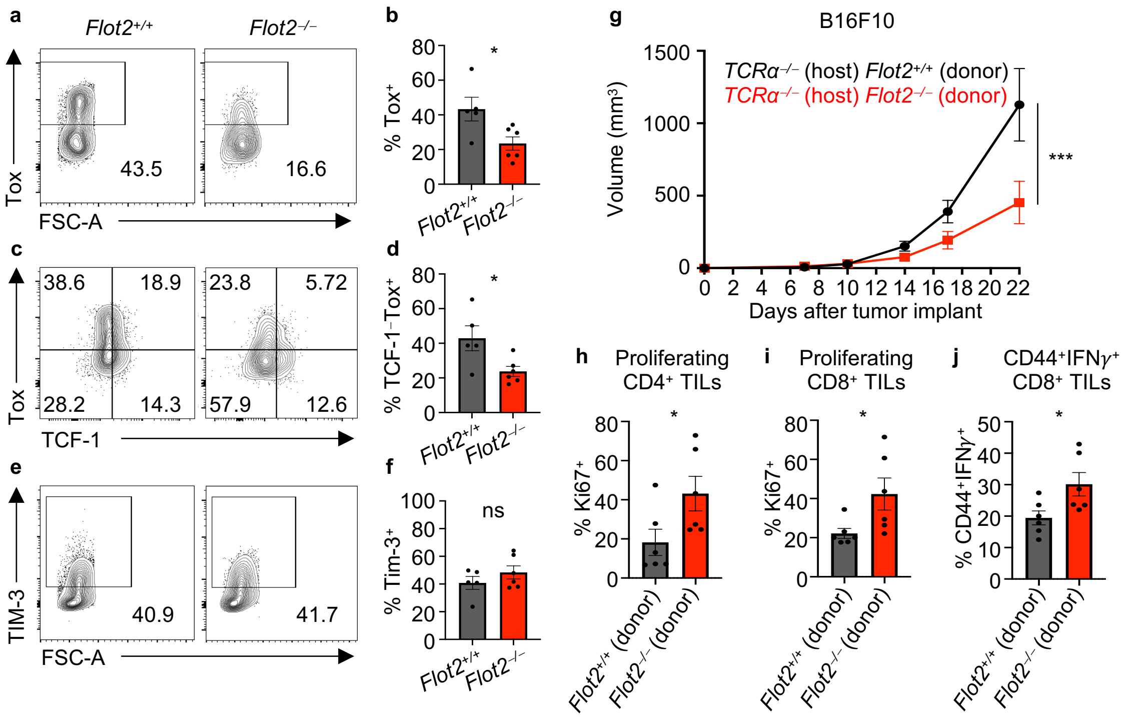


**Supplementary Figure 2. Flot2 deficiency reduces T cell functional exhaustion and enhances anti-tumor T cell responses.** **a**-**f** Flow cytometric analysis of TILs in MC38 tumor-bearing *Flot2*$^{+/+}$ or *Flot2*$^{-/-}$ mice: Representative plots (**a**, **c**, **e**) are shown. Tox$^{+}$ (**b**), TCF-1$^{-}$Tox$^{+}$ (**d**), and Tim-3$^{+}$ (**f**) populations within 7AAD$^{-}$CD45.2$^{+}$TCRβ$^{+}$CD8$^{+}$ population are depicted. **g** B16F10 tumor volume in mixed bone marrow chimeras reconstituted with a 1:5 ratio mixture of bone marrow cells from either *Flot2*$^{+/+}$ or *Flot2*$^{-/-}$ mice and *TCRα*$^{-/-}$ mice (n = 8 per group). **h-j** Flow cytometric analysis of TILs. Ki67$^{+}$ populations among 7AAD$^{-}$CD45$^{+}$CD4$^{+}$ population (**h**) or 7AAD$^{-}$CD45$^{+}$CD8$^{+}$ population (**i**), and CD44$^{+}$IFN$\gamma^{+}$ populations within 7AAD$^{-}$CD45$^{+}$CD8$^{+}$ population (**j**) are shown. Data are representative of two independent experiments (**a**-**j**). Data were analyzed by unpaired t-test (**b**, **d**, **f**, **h-j**) or two-way ANOVA (**g**). Error bars denote SEM; *P<0.05; ***P<0.001. ns = non-significant.

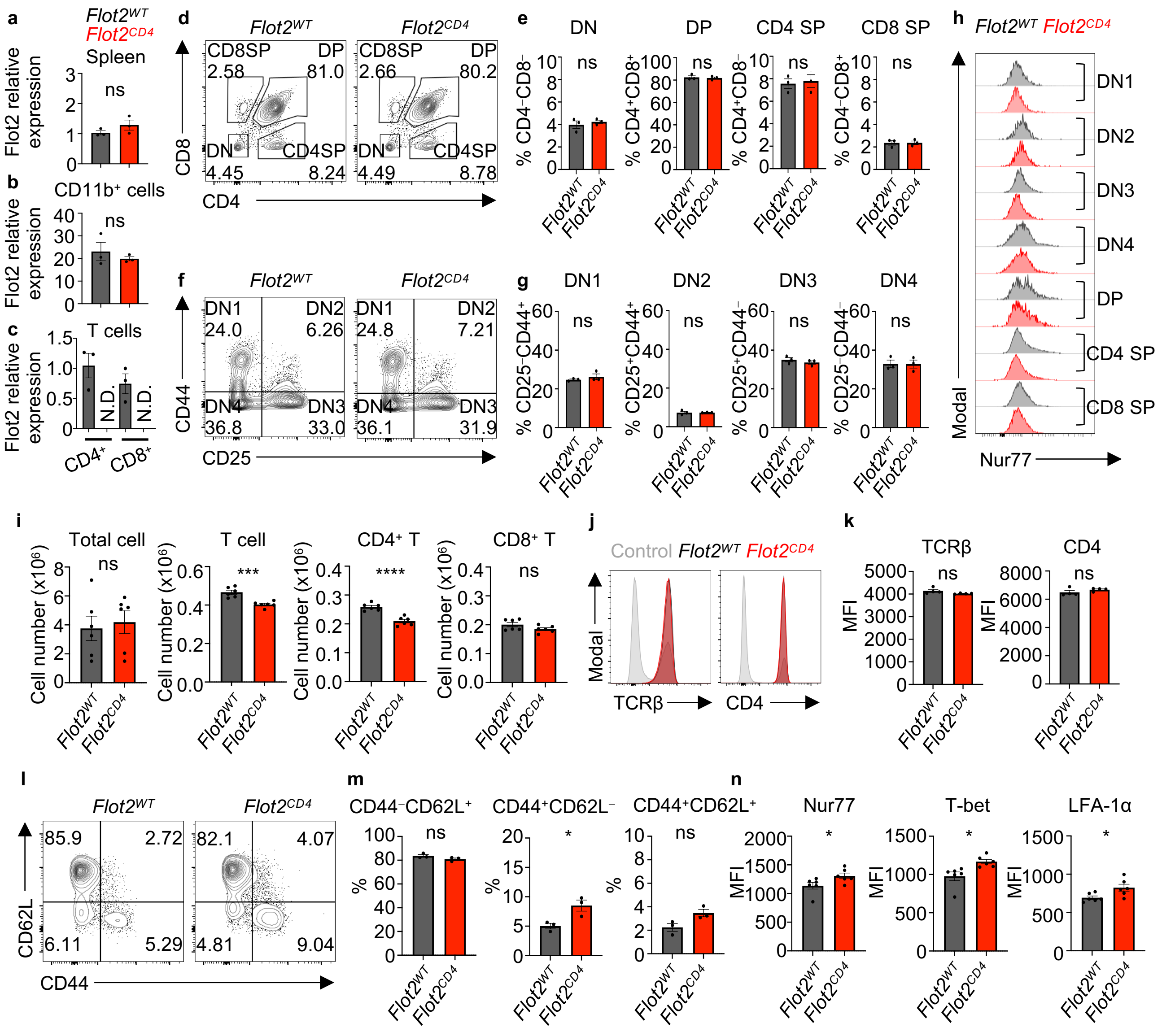


**Supplementary Figure 3. Steady state analysis of *Flot2*$^{WT}$ and *Flot2*$^{CD4}$.**

**a**-**c** Flot2 mRNA expression was comparable in total spleen (**a**) or CD11b$^+$ cells (**b**) but was fully deleted in both CD4$^+$ and CD8$^+$ T cells of *Flot2*$^{CD4}$ mice (**c**). **d**-**h** Flow cytometric analysis of thymocytes in *Flot2*$^{WT}$ and *Flot2*$^{CD4}$ mice at steady state. Representative plots (**d**, **f**) are shown. CD4$^-$CD8$^-$ (DN), CD4$^+$CD8$^+$ (DP), CD4$^+$CD8$^-$ (CD4 SP), and CD4$^-$CD8$^+$ (CD8 SP) populations among singlet thymocytes (**e**) and CD25$^-$CD44$^+$ (DN1), CD25$^+$CD44$^+$ (DN2), CD25$^+$CD44$^-$ (DN3), CD25$^-$CD44$^-$ (DN4) populations within the DN subset (**g**) in steady state *Flot2*$^{WT}$ or *Flot2*$^{CD4}$ mice are presented. Flow cytometric histogram plot of Nur77 expression at each stage of thymocytes are displayed (**h**). **i** Cell numbers of total cells, T cells, CD4$^+$ T cells, CD8$^+$ T cells in inguinal lymph nodes of steady state *Flot2*$^{WT}$ or *Flot2*$^{CD4}$ mice. **j**, **k** Flow cytometric analysis of TCRβ and CD4 expression on viable lymphocytes at steady state. Representative plots (**j**) and mean fluorescence intensity (MFI) quantification (**k**) are shown. **l**-**n** Flow cytometric analysis of naïve, effector, central memory populations and Nur77, T-bet, LFA-1α expression within steady state lymph node CD4$^+$ T cells. Representative plots of each populations are shown (**l**). CD44$^-$CD62L$^+$ (naive), CD44$^+$CD62L$^-$ (effector), CD44$^+$CD62L$^+$ (central memory) populations (**m**) and MFI quantification of Nur77, T-bet, LFA-1α (**n**) are provided. Data are representative of two independent experiments (**a**-**n**). Data were analyzed by unpaired t-test (**a**-**c**, **e**, **g**, **i**, **k**, **m**, **n**). Error bars denote SEM; *P<0.05; ***P<0.001; ****P<0.0001. ns = non-significant. N.D. = not detected.

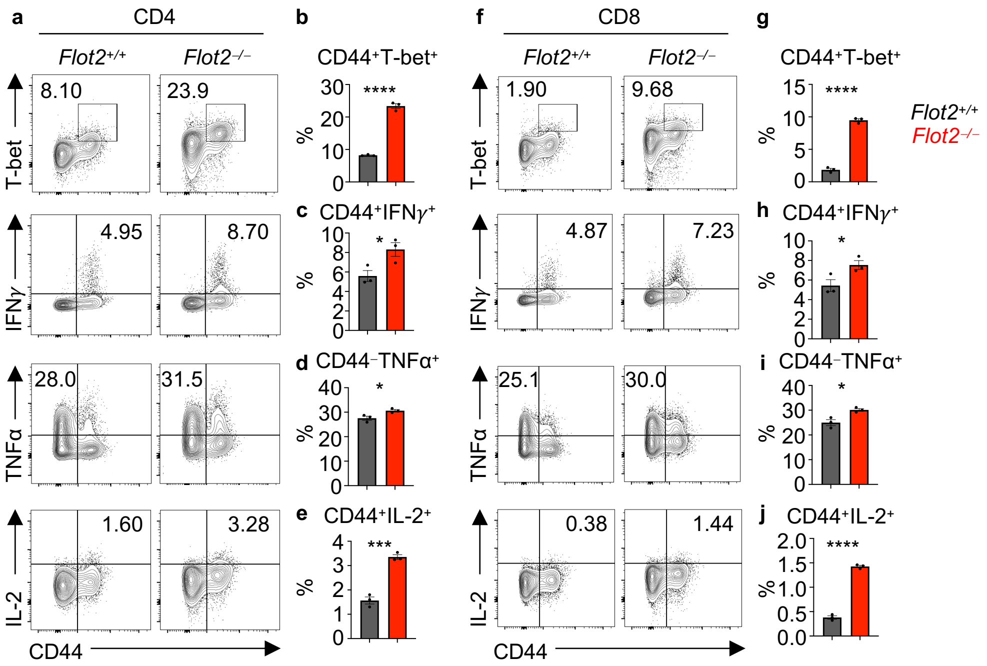


**Supplementary Figure 4. Flot2 deletion boosts effector T cell responses in Listeria monocytogenes-infected mice.**
**a**-**j** Flow cytometric analysis of splenic T cells from *Listeria*-infected *Flot2*$^{+/+}$ or *Flot2*$^{-/-}$ mice: Representative plots (**a**, **f**) are shown. CD44$^{+}$T-bet$^{+}$ (**b**), CD44$^{+}$IFN$\gamma^{+}$ (**c**), CD44$^{-}$TNFα$^{+}$ (**d**), and CD44$^{+}$IL-2$^{+}$ (**e**) populations within viable CD45$^{+}$TCRβ$^{+}$CD4$^{+}$ population, and CD44$^{+}$T-bet$^{+}$ (**g**), CD44$^{+}$IFN$\gamma^{+}$ (**h**), CD44$^{-}$TNFα$^{+}$ (**i**), and CD44$^{+}$IL-2$^{+}$ (**j**) populations within viable CD45$^{+}$TCRβ$^{+}$CD8$^{+}$ population are shown. Data are representative of two independent experiments (**a**-**j**). Data were analyzed by unpaired t-test (**b-e, g-j**). Error bars denote SEM; *P<0.05; ***P<0.001; ****P<0.0001.

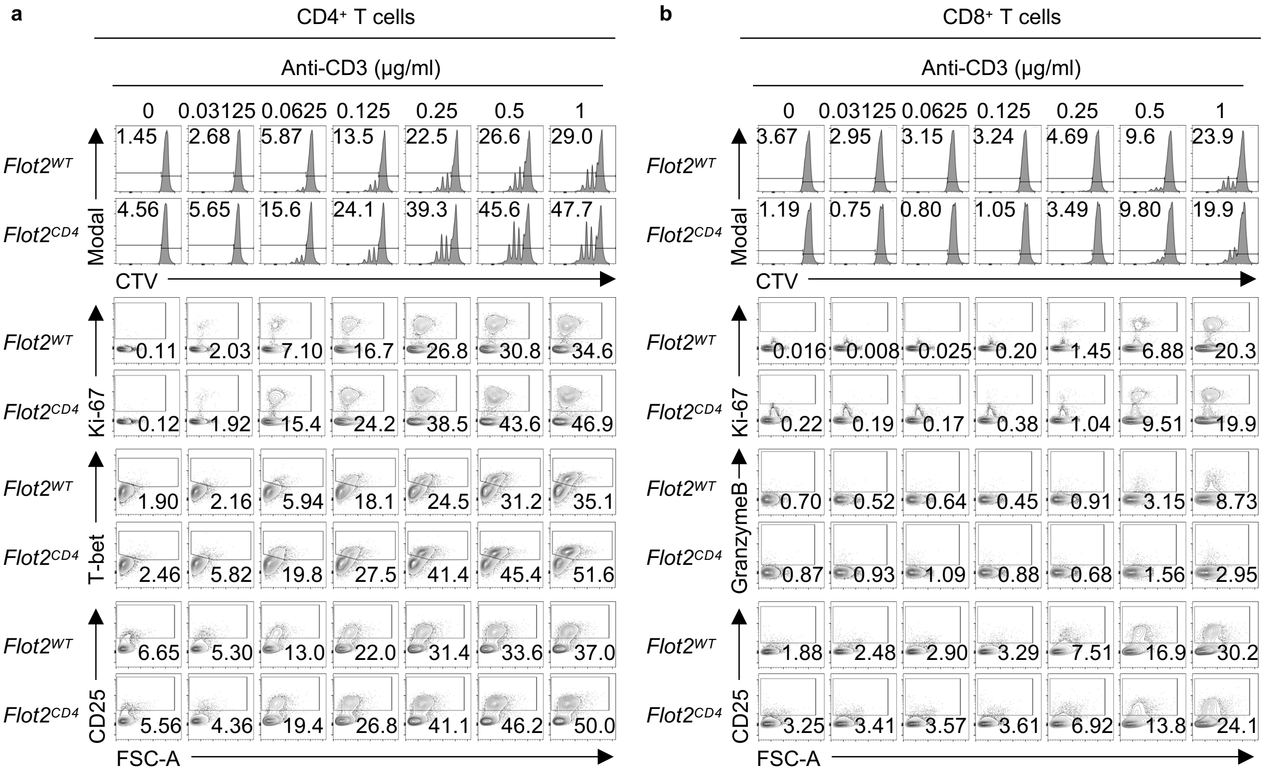


**Supplementary Figure 5. *Flot2$^{CD4}$* CD4⁺ but not CD8⁺ T cells displayed enhanced responses to *in vitro* stimulation.**
Representative plots of flow cytometric analysis of naïve CD4⁺ (**a**) or CD8⁺ T cells (**b**) that were stimulated *in vitro* for 72 hours with varying doses of plate-bound anti-CD3, alongside a fixed dose of soluble anti-CD28 (1 µg/ml). CTV⁻, Ki67⁺, T-bet⁺, and CD25⁺ populations within viable TCRβ⁺CD4⁺ population (**a**) or CTV⁻, Ki67⁺, Granzyme B⁺, and CD25⁺ populations within viable TCRβ⁺CD8⁺ population (**b**) are shown. Data are representative of two independent experiments (**a**, **b**).

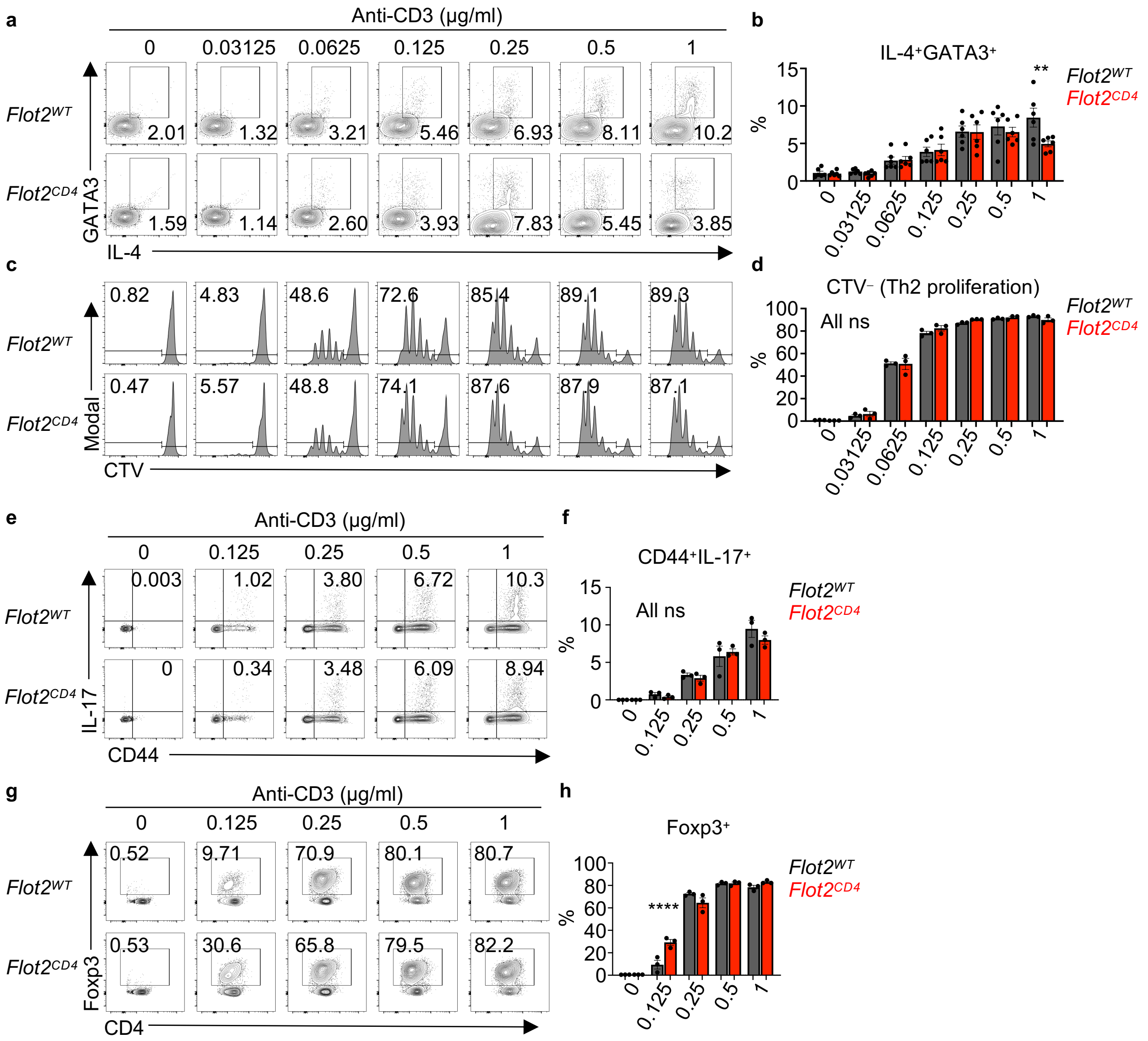


**Supplementary Figure 6. Flot2 ablation does not affect Th2 and Th17 differentiation, while enhancing Treg differentiation at weak TCR stimulation.**

**a-h** Naïve CD4+ T cells were purified and differentiated towards the Th2 (**a**-**d**), Th17 (**e**, **f**) and Treg (**g**, **h**) subtypes *in vitro*, followed by flow cytometric analysis. Representative plots (**a**, **c**, **e**, **g**) are shown. IL-4+GATA3+ (**b**), CTV− (**d**), CD44+IL-17+ (**f**), and Foxp3+ (**h**) populations within viable TCRβ+CD4+ population are shown. Data are representative of two independent experiments (**a**-**h**). Data were analyzed by one-way ANOVA followed with Sidak's multiple comparison tests (**b**, **d**, **f**, **h**). Error bars denote SEM; **P<0.01; ****P<0.0001. ns = non-significant.

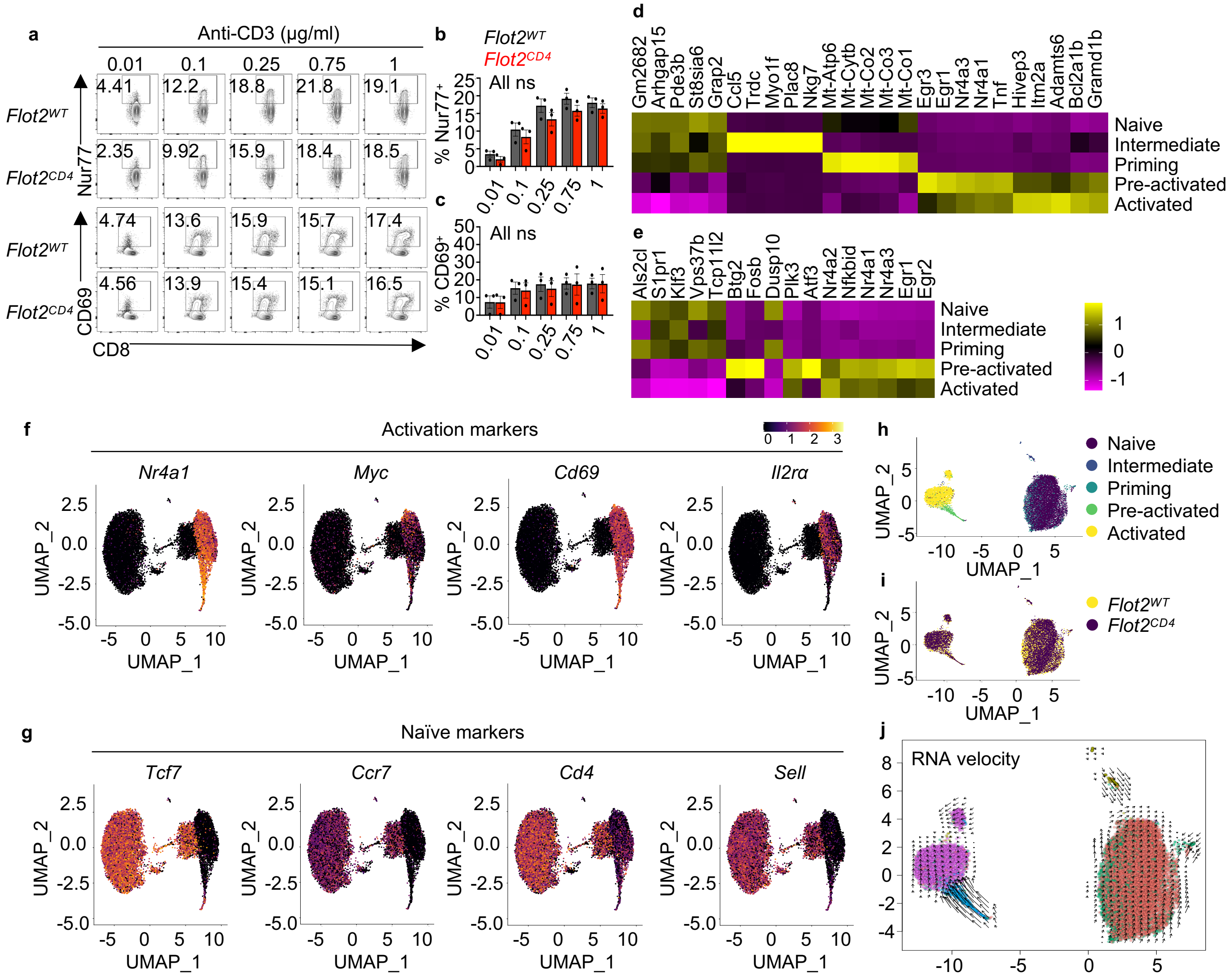


**Supplementary Figure 7. Flot2 ablation does not impact the early activation of CD8$^+$ T cells upon *in vitro* stimulation, nor does it affect the transcriptional activation trajectory of CD4$^+$ T cells.**

**a-c** Naïve CD8$^+$ T cells were purified and stimulated *in vitro* for 3 hours (**b**) or 24 hours (**c**) with varying doses of plate-bound anti-CD3, alongside a fixed dose of soluble anti-CD28 (1 μg/ml), followed by flow cytometric analysis to assess TCR signaling (Nur77) and early T cell activation (CD69). Representative plots (**a**) are shown. Nur77$^+$ (**b**) and CD69$^+$ (**c**) populations within viable TCRβ$^+$CD8$^+$ population are indicated. **d**, **e** Gene expression heatmap from scRNA-seq analysis. Top markers for clustering (**d**) and gene set related to early T cell activation (**e**) are shown. **f**, **g** Expression of T cell activation (**f**) or naïve state (**g**) marker genes over the UMAP dot plots. **h-j** T cell activation trajectory fitted by RNA velocity analysis. T cell functional clustering (**h**), genotype (**i**), and RNA velocity (**j**) results are shown. Data are pooled from three independent experiments and were analyzed by one-way ANOVA followed with Sidak's multiple comparison tests (**a-c**). Error bars denote SEM. ns = non-significant.

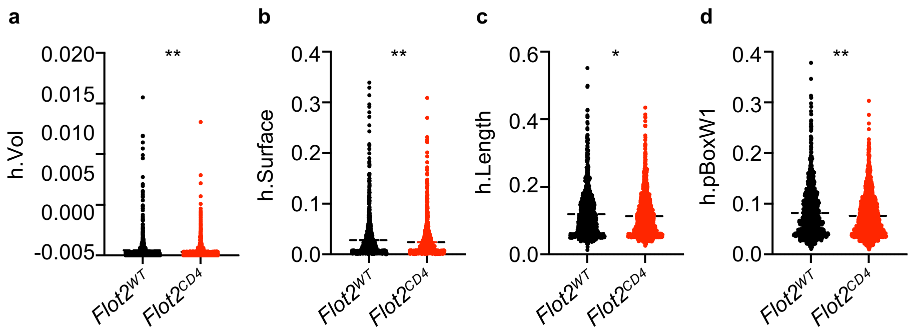


**Supplementary Figure 8. Convex hull geometry analysis of TCR nanoclusters.**
**a**-**d** Convex hull geometry analysis of TCRβ⁺ nanoclusters in *Flot2*$^{WT}$ and *Flot2*$^{CD4}$ naïve CD4⁺ T cells: Volume enclosed by the convex hull of the cluster in cubic micrometers (h.Vol; **a**), surface of the convex hull of the cluster in squared micrometers (h.Surface; **b**), the largest length of the convex hull of the cluster in micrometers (h.Length; **c**), and the largest width of the convex hull principal box perpendicular to the convex hull length in micrometers (h.pBoxW1; **d**) are quantified and displayed. Data are representative of two independent experiments (**a**-**d**). Data were analyzed by unpaired t-test (**a**-**d**). Error bars denote SEM; *P<0.05; **P<0.01.

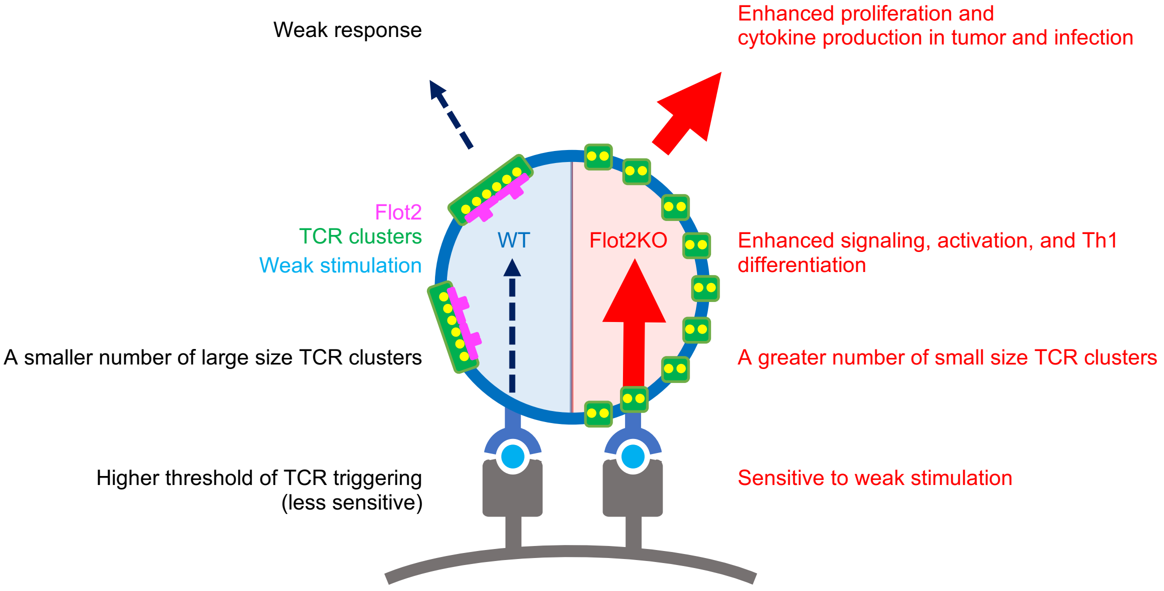


**Supplementary Figure 9. Flotillin-2 dampens T cell antigen-sensitivity and functionality.**
